# StormGraph: A graph-based algorithm for quantitative clustering analysis of diverse single-molecule localization microscopy data

**DOI:** 10.1101/515627

**Authors:** Joshua M. Scurll, Libin Abraham, Da Wei Zheng, Reza Tafteh, Keng C. Chou, Michael R. Gold, Daniel Coombs

## Abstract

Clustering of proteins is crucial for many cellular processes and can be imaged at nanoscale resolution using single-molecule localization microscopy (SMLM). Ideally, molecular clustering in regions of interest (ROIs) from SMLM images would be assessed using computational methods that are robust to sample and experimental heterogeneity, account for uncertainties in localization data, can analyze both 2D and 3D data, and have practical computational requirements in terms of time and hardware. While analyzing surface protein clustering on B lymphocytes using SMLM, we encountered limitations with existing cluster analysis methods. This inspired us to develop StormGraph, an algorithm using graph theory and community detection to identify clusters in heterogeneous sets of 2D and 3D SMLM data while accounting for localization uncertainties. StormGraph generates both multi-level and single-level clusterings and can quantify cluster overlap for two-color SMLM data. Importantly, StormGraph automatically determines scale-dependent thresholds from the data using scale-independent input parameters. This makes identical choices of input parameter values suitable for disparate ROIs, eliminating the need to tune parameters for different ROIs in heterogeneous SMLM datasets. We show that StormGraph outperforms existing algorithms at analyzing heterogeneous sets of simulated SMLM ROIs where ground-truth clusters are known. Applying StormGraph to real SMLM data in 2D, we reveal that B-cell antigen receptors (BCRs) reside in a heterogeneous combination of small and large clusters following stimulation, which suggests for the first time that two conflicting models of BCR activation are not mutually exclusive. We also demonstrate application of StormGraph to real two-color and 3D SMLM data.

## Introduction

Single-molecule localization microscopy (SMLM) is commonly used to investigate nanoscale clustering of cell-membrane and intracellular proteins in selected cellular regions of interest (ROIs) (1; 2; 3; 4; 5; 6; 7; 8; 9; 10). SMLM techniques, such as direct stochastic optical reconstruction microscopy (dSTORM) (11; 12) and photoactivated localization microscopy (PALM) (13), overcome the diffraction limit of conventional microscopy by acquiring many sequential images, each containing very few fluorescing labels. Individual labels can then be computationally super-resolved and precisely localized to generate localization coordinates, often with estimated positional uncertainties (14; 15; 16). This is possible in both two and three dimensions (17; 18; 19; 20).

SMLM data usually exhibits ROI-to-ROI and within-ROI heterogeneity due to biological and technical variability between imaged cells and to the spatial heterogeneity of the plasma membrane, intracellular compartments, and cytoplasm. Nevertheless, clustering is frequently analyzed using spatial summary statistics that fail to capture the heterogeneity of clusters within ROIs, for example Ripley’s functions (21; 22). Over the last six years, we have regularly applied such methods ourselves to dSTORM data as part of investigations of the spatial distributions of cell-surface molecules on B cells in relation to B-cell antigen receptor (BCR) signaling. For more informative analysis, clusters can be individually quantified by using a clustering algorithm to assign localizations to specific clusters (23; 24; 25; 26; 27; 28; 29; 30; 31). However, challenged by large numbers and diversity of ROIs, we had difficulty using existing algorithms to achieve consistent analysis of nanoscale BCR clustering on tens or hundreds of B cells imaged by dSTORM. We consequently developed our own method, which we call StormGraph.

Herein, we describe and demonstrate the use of StormGraph, a comprehensive graph-based clustering algorithm inspired by PhenoGraph (32) from the single-cell cytometry field. StormGraph converts SMLM data into a graph using localization coordinates and their uncertainties to specify nodes and weighted edges. It then utilizes graph theory and community detection (33) to assign nodes to clusters. StormGraph makes no assumptions about the shapes of clusters, has both 2D and 3D implementations, and can quantify cluster overlap for two-color SMLM data. Using Monte Carlo simulations, StormGraph automatically adapts key thresholds to each ROI independently. This allows users to keep input parameters fixed across all ROIs in an experiment without introducing systematic bias to the analysis. Additionally, StormGraph generates a multi-level (i.e. hierarchical) clustering whereby clusters are recursively composed of smaller clusters. Examples of multi-level clustering possible in SMLM data include multiscale organization of RNA polymerase in *Escherichia coli* (34) and clustering of receptor oligomers, themselves clusters of molecules, into lipid rafts. By outputting a multi-level clustering, StormGraph simplifies multiscale cluster analysis compared to repeatedly changing parameters for existing methods. Notwithstanding, StormGraph also generates an appropriate single-level clustering to facilitate easy-to-interpret analysis. To streamline analysis of ROIs from many samples, we developed software to manually crop ROIs from one or two SMLM color channels and subsequently batch analyze them with StormGraph. We believe that StormGraph has certain advantages over other clustering algorithms in the SMLM literature.

The most widely used clustering algorithms in SMLM literature, including Density-Based Spatial Clustering of Applications with Noise (DBSCAN) (23), identify clusters based on a user-specified minimum number of points within a user-specified radius. The optimal values of their parameters depend on the density of the data, but the localization density typically varies between ROIs. Consequently, the common practice of fixing these parameters while analyzing multiple ROIs can systematically bias cluster analyses because the parameter choice may be inappropriate for many of the ROIs. The alternative approach of choosing different parameter values for each ROI would introduce an enormous amount of subjective bias, especially because parameter selection is usually challenging even for a single ROI. Alternative algorithms based on Voronoi diagrams have been developed for 2D (24; 25) and 3D SMLM data (26) and address these problems in different ways.

A Voronoi diagram divides an ROI into tessellated “Voronoi cells” (polygons in 2D or polyhedra in 3D) whereby each Voronoi cell encloses one localization and all regions of space that are closer to it than to any other localization. Both SR-Tesseler (24) and ClusterViSu (25) construct Voronoi diagrams, then apply thresholds to the Voronoi cells, and finally group adjacent Voronoi cells into clusters. However, they differ in how they determine thresholds. SR-Tesseler provides users with several options for setting thresholds, but the leading option sets a single threshold on density (defined as reciprocal of Voronoi polygon area) equal to a user-specified constant multiplicative factor, *α*, of the average density of localizations in the ROI. This automatically adapts the density threshold to the average localization density in each ROI but neglects the variance that would be expected for randomly distributed localizations. ClusterViSu similarly applies a single threshold to Voronoi polygon areas, but it uses Monte Carlo simulations to automatically set the density threshold equal to the cutoff at which Voronoi polygon areas appear more frequently in the actual data than in random data. This is in contrast to the spatially uniform (i.e. evenly spaced) null reference distribution of localizations that SR-Tesseler uses. Although the approach of ClusterViSu adds computational expense, it eliminates user influence from calculation of the threshold, and the authors of ClusterViSu demonstrated that it is more robust than SR-Tesseler over a large range of background localization densities (25). Inspired by ClusterViSu, StormGraph overcomes the parameter selection problem of density-based clustering algorithms (e.g. DBSCAN) by using density-independent input parameters, which users can keep fixed, and then automatically adapting density-dependent thresholds to each ROI using Monte Carlo simulations.

Although ClusterViSu is an attractive method for analyzing datasets with ROI-to-ROI heterogeneity, none of the methods described so far account for positional uncertainties in SMLM localizations. These are often output from the initial processing of raw data, alongside the most probable localization positions, and provide additional information that can be exploited to improve clustering results. Two methods based on DBSCAN use this information to some extent. One is a pixelated method specifically for 2D data (27), and the other corrects cluster-size distributions, but not actual clusters, determined using regular DBSCAN (28). Most appealing, a Bayesian, model-based clustering algorithm applicable to 2D or 3D data (29; 30) builds the positional uncertainty of each localization into the cluster detection process. However, it assumes that all clusters in an ROI have circular or spherical Gaussian profiles of similar size. This potentially limits its suitability for data with clusters of elongated or unusual shapes or heterogeneous sizes. Furthermore, among existing SMLM cluster analysis methods, the Bayesian method has the most user-adjustable settings, which can be non-intuitive (e.g. Dirichlet concentration parameter) or difficult to determine (e.g. Bayesian priors). Also, slow computation times limit its practicality, with the method typically requiring ~30 minutes on a standard desktop computer to analyze one ROI containing 1,000 localizations (30), which is less than one tenth of the number of localizations that we routinely acquire per ROI. In contrast, StormGraph makes use of all available positional uncertainties for 2D or 3D localization data without imposing assumptions about cluster shapes or sizes or requiring excessive computation times.

Herein, we describe StormGraph and its capabilities, and we use simulated data to compare its performance to that of DBSCAN, ClusterViSu, and the Bayesian method. We then apply StormGraph to characterize nanoscale BCR clustering from heterogeneous 2D SMLM data, which delivers novel insights into BCR organization in resting and activated B cells. We also demonstrate StormGraph’s ability to quantify 3D clusters of the lysosomal protein LAMP-1 and to quantify cluster overlap for two-color SMLM data.

## Results

### The StormGraph algorithm

As input to the clustering algorithm, StormGraph takes ROIs that have been selected manually from field-of-view SMLM images, for example using our software. The ROIs should be completely enclosed within the boundaries of imaged cells because StormGraph, like other cluster analysis methods including ClusterViSu and Ripley’s functions, compares the data to a completely uniformly random ‘null’ distribution of points. This choice of null distribution is not appropriate for any ROI that contains unoccupied coverslip space as such spaces should remain empty in the null distribution. Besides, keeping ROIs within cell boundaries is generally necessary for 2D data to avoid artifacts of projecting 3D cell-membrane curvature onto two dimensions.

To identify clusters in an ROI, dense localization neighborhoods must be identified. To this end, StormGraph first determines an ROI-specific length scale *r*_0_ from the data using either of two methods (see Methods and Figure S1). The preferred method uses an input parameter *k*, which specifies a number of nearest neighbors of each localization. The computation of *r*_0_ then resembles the automatic threshold computation in ClusterViSu but using distance to *k*^th^ nearest neighbor (kNN) in lieu of Voronoi polygon area. To reduce user input, StormGraph alternatively offers a fully automatic but heuristic method to compute *r*_0_ by seeking a balance between inter-localization and inter-cluster distances without any user-adjustable parameters. This heuristic method is intended for visually well clustered data with very few dispersed localizations between clusters, but we nonetheless found that it produces comparable StormGraph results to the universally applicable kNN method even for unintended use cases.

Next, using the localizations as nodes (Figure 1a), StormGraph essentially constructs a weighted *r*_0_-neighborhood graph (Figure 1b) as follows. Define the similarity, *s_ij_*, of two nodes, *i* and *j,* to be

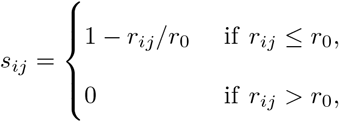

where *r_ij_* is the Euclidean distance between nodes *i* and *j*. If localization coordinate uncertainties are unknown, then StormGraph assigns to each node pair {*i,j*} an edge of weight *W_ij_* = *s_ij_*. Otherwise, StormGraph uses the uncertainties to estimate 〈*s_ij_*〉, the expectation of the similarity *s_ij_*, from Monte Carlo simulations (Methods) and assigns *W_ij_* = 〈*s_ij_*〉.

**Figure 1:**
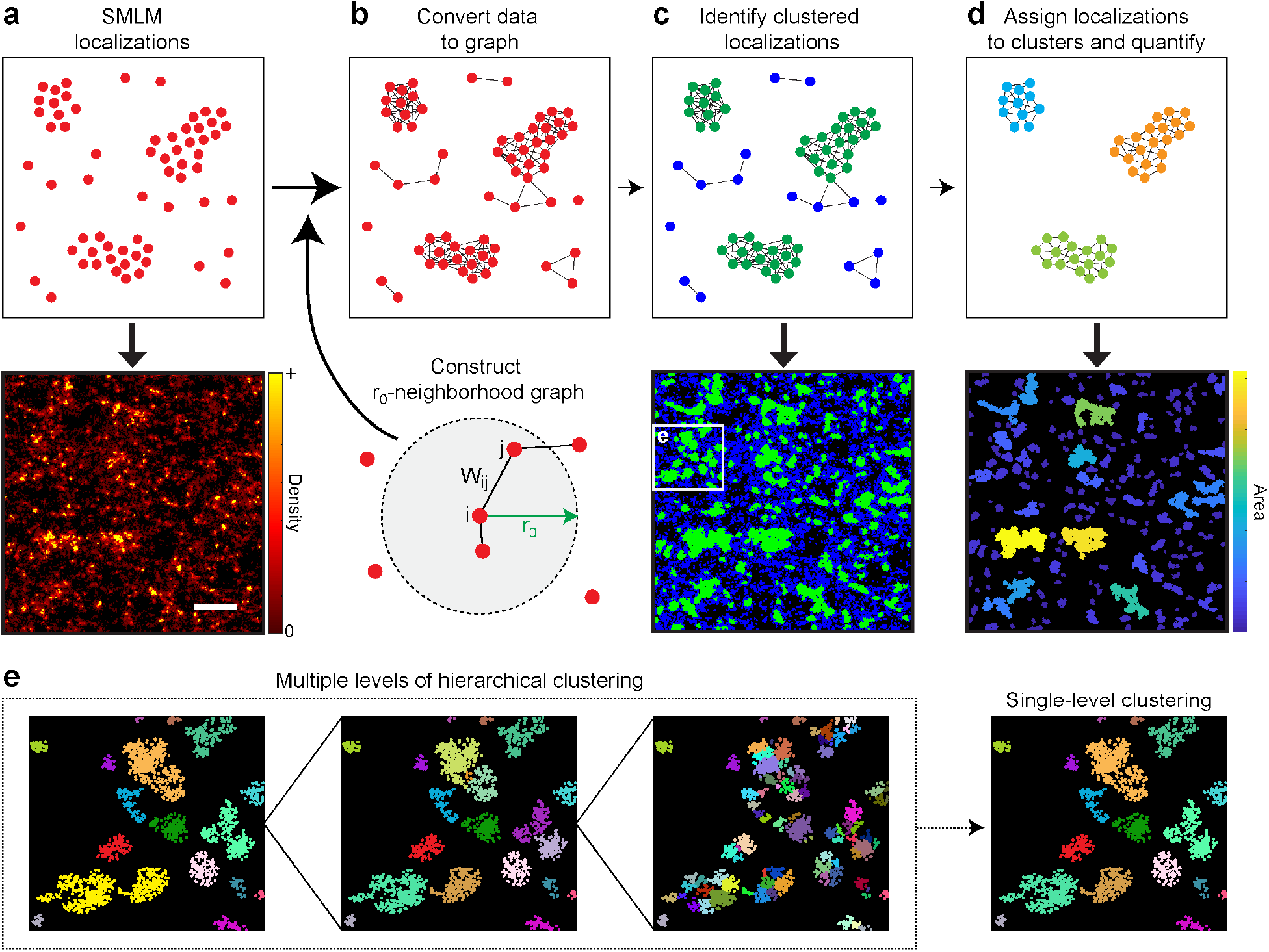
Schematic illustration of StormGraph’s workflow. **(a–d)** SMLM localizations are used as nodes (a) and converted into a weighted graph (b). This graph is based on *r*_0_-neighborhood graphs, where edges connect each node to all other nodes within a distance *r*_0_. Two nodes i and *j* are connected by an edge of weight *W_ij_*, which describes their similarity based on the distance between them and, if known, their positional uncertainties. Nodes are classified as either clustered (green) or unclustered (blue) based on their node degree, i.e. sum of adjacent edge weights, (c). A new graph is constructed from only the clustered nodes, which are then assigned to specific clusters using a community detection algorithm (d). Cluster properties (e.g. area) can then be quantified. The bottom panels in (a), (c), and (d) illustrate each step for an actual SMLM region of interest (scale bar = 500 nm). **(e)** StormGraph identifies a hierarchy of clusters at multiple scales and then additionally generates a single-level clustering from the hierarchy. Shown are three different levels from the cluster hierarchy for the region in the white box in the lower panel of (c), along with the single-level clustering for this region. Colors distinguish different clusters.

At this stage, unclustered localizations are identified and removed by applying a threshold to the weighted node degree,

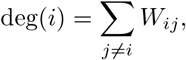

a proxy for local density. In principle, nodes can be classified as unclustered and removed (reported as ‘cluster 0’ in StormGraph’s output) if their degree falls below a data-dependent threshold (Figure 1c). StormGraph automatically determines this threshold from random point clouds using the user-defined parameter *α* (Methods), which is a per-localization significance level for the null hypothesis that localizations are randomly distributed. StormGraph’s default value of *α* is 0.05, but users may alter this within the range 0 ≤ *α* ≤ 1. Figure S2 illustrates the effects of varying the parameters *α* and *k* for an SMLM ROI with ambiguous clusters.

The graph is then regenerated using a new *r*_0_ value determined completely automatically, using the heuristic method, from only the retained nodes. If localization coordinate uncertainties are available, edges are subsequently pruned from the graph to ensure that all retained pairs of edges have at least an estimated 50% probability of co-occurring in the r_0_-neighborhood graph for the unknown true localization positions (Methods). Without edge pruning, the average graph contains all edges that occur in at least one Monte Carlo simulation. Consequently, the average graph can contain combinations of edges that rarely or never co-occur in the simulations and therefore its connectivity will not necessarily reflect the true cluster structure of the data. The edge pruning addresses this by ensuring that any pair of retained edges co-occur at least as often as not. StormGraph then finds a hierarchy of node clusters (Figure 1e) using the multi-level Infomap community detection algorithm (35), followed by additional cluster merging when warranted (Methods).

To obtain a single-level clustering from the hierarchy, we developed a novel, fast method motivated by the idea of consensus clustering (36; 37). Briefly, starting at the top of the cluster hierarchy, clusters are recursively divided into their coarsest constituent subclusters until they no longer bear similarity to connected components of an alternative neighborhood graph based on mutual nearest neighbors (Methods). Optionally, the user may specify a desired minimum number of localizations per cluster (MinCluSize). This will impose a limit on how far clusters can be subdivided if the stopping criterion is not already met and it will exclude clusters containing fewer than MinCluSize localizations from downstream analyses. As output, StormGraph provides the single-level and multi-level cluster assignments of every localization. Combined with localization coordinates, this provides the information necessary to quantify individual cluster properties, such as area (Figure 1d) and number of localizations per cluster. Our software automatically quantifies the single-level and coarsest-level clusterings.

A common caveat of SMLM is multiple counting of single molecules, often causing single molecules to spuriously appear as clusters. This can be due to multiple labeling of single molecules or to repeated photoblinking of individual fluorophores. Therefore, StormGraph includes optional functionality, using a statistical approach revolving around localization uncertainties, to reclassify as unclustered localizations any putative clusters that cannot be confidently distinguished from multiply counted single molecules (Methods). However, like other clustering algorithms, StormGraph does not attempt to infer the number or positions of actual molecules. Hence, we caution that reporting localization numbers, instead of cluster areas for example, can mislead biological interpretation of real SMLM data.

We note one potential limitation of StormGraph. Infomap, like other leading community detection algorithms, is not guaranteed to produce identical results from repeat runs. Monte Carlo simulations also introduce stochastic variability between repeat runs of StormGraph. However, discrepancies between the clusters output by identical repeat runs of StormGraph indicate that clusters have ambiguous boundaries and therefore that there are multiple ways in which the localizations can be rationally partitioned into clusters. Nonetheless, we tested the reproducibility of StormGraph and found that the clusters generated by identical repeat runs of StormGraph for a heterogeneous dSTORM ROI containing visually ill-defined clusters were highly similar (Figure S3; see Methods).

### Validation using simulated data and comparison to other algorithms

To compare StormGraph with DBSCAN and ClusterViSu, we simulated 64 diverse 2 μm × 2 μm ROIs containing isolated and heterogeneously aggregated circular nanoclusters (e.g. Figure 2a–c; Methods). Outside the clusters we added randomly distributed molecules. Individual simulated molecules were allowed to yield multiple localizations, each with a positional uncertainty sampled from a real dSTORM experiment. We tested both the automatic and kNN (*k* = 10, 15 or 20) methods for determining *r*_0_ while maintaining *α* = 0.05. Although the MinCluSize parameter is not required by StormGraph, we found that ClusterViSu would often detect many small, spurious ‘clusters’, even as small as just one localization, if we did not set a minimum number of localizations needed for a cluster to be retained. We therefore set a minimum cluster size of 5 localizations in ClusterViSu, and since MinCluSize functions similarly in StormGraph, we set it equally in order to make a fair comparison between the two algorithms. For DBSCAN, we tested 16 different parameter choices based on the underlying parameters used for data simulation, although such knowledge is generally unavailable for real data. To assess cluster assignments by each algorithm, we used normalized mutual information (NMI) (38) and mean F-measure (39). Higher values indicate superior performance. We also evaluated resulting errors in cluster quantification. For each simulated ROI, we calculated the resultant errors in the mean (*μ*) and standard deviation (*σ*) of the number of localizations per cluster and in the overall percentage of localizations assigned to clusters. We report these errors as fractions of (i.e. relative to) the ground-truth quantification values. Thus, errors lie between −1 (i.e. −100% error) and +∞ with 0 indicating zero error.

**Figure 2:**
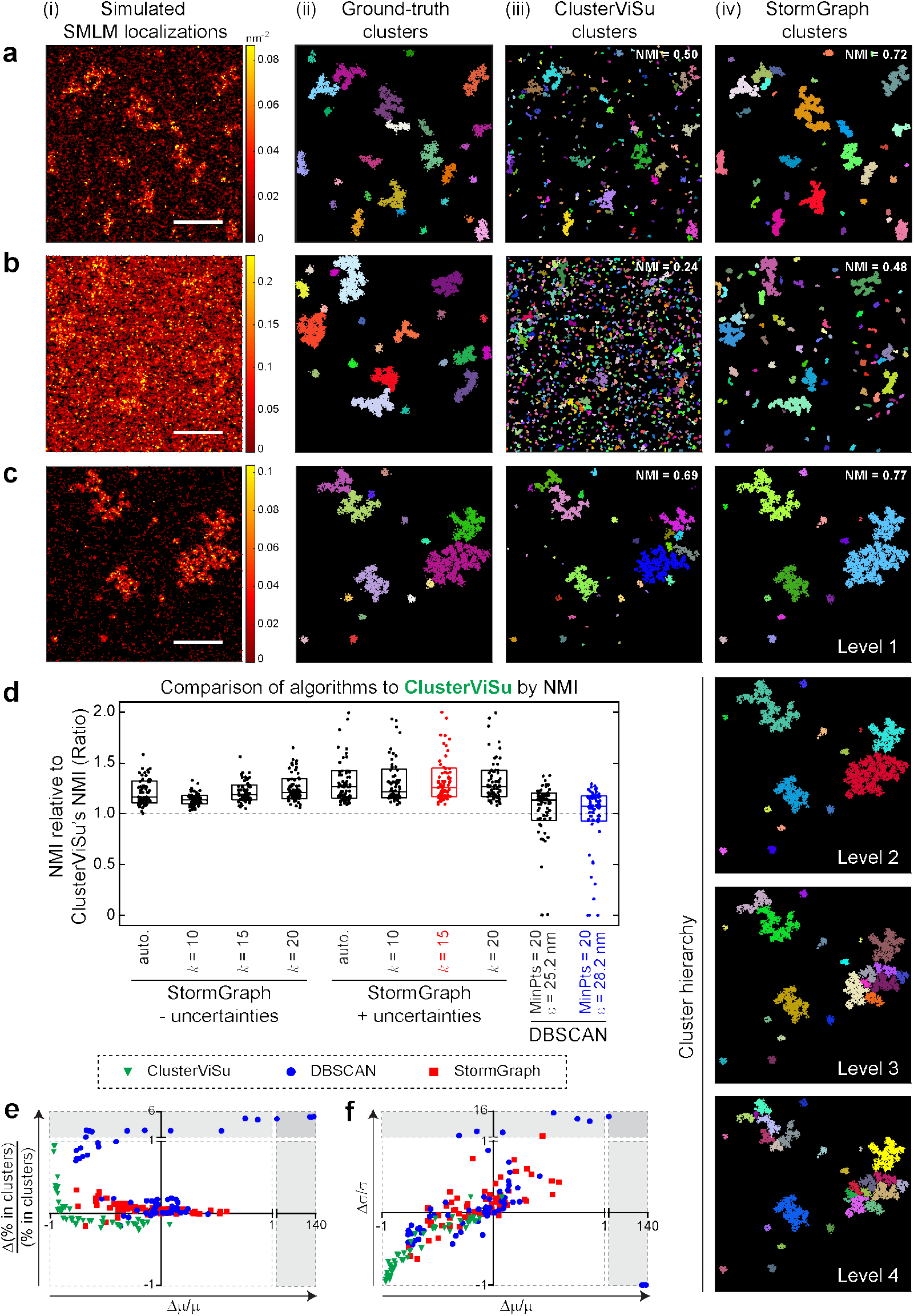
StormGraph consistently outperforms ClusterViSu and DBSCAN on simulated data. **(a–c)** (i) Examples of simulated data (color bar = density, scale bar = 500 nm), (ii) their ground-truth clusters, and cluster assignment results with corresponding absolute normalized mutual information (NMI) values (1 = perfect) for (iii) ClusterViSu and (iv) StormGraph (+ uncertainties; *k* = 15; single-level). Colors distinguish distinct clusters. Also shown are four levels (Levels 1-4) of the multi-level cluster hierarchy output by StormGraph for (c)(i). In this particular example, the single-level clustering (c)(iv) generated by StormGraph and the coarsest level (Level 1) of the cluster hierarchy were identical, but this is not the case in general. **(d)** Ratios of the performances of StormGraph and DBSCAN to the performances of ClusterViSu when performance was evaluated by NMI of cluster assignments compared to ground truth. Ratios > 1 (respectively < 1) indicate that StormGraph or DBSCAN performed better (respectively worse) than ClusterViSu. A total of 64 simulated ROIs were analyzed, including the ones shown in (a-c). We terminated ClusterViSu if it took longer than 2 h to analyze an ROI, which resulted in the exclusion of 15 out of 64 ROIs. StormGraph was run either with (+) or without (-) localization uncertainties and using either the heuristic method (auto.) or the kNN method with *k* = 10, 15, or 20 to set *r*_0_. DBSCAN was implemented using 16 different selections of its two parameters, MinPts and *ε*, of which the two bestperforming are shown. Cluster assignment results were scored using NMI and the scores for StormGraph and DBSCAN were divided by the scores for ClusterViSu. Each dot in the figure shows the NMI ratio for one of the 49 simulated ROIs analyzed by all three algorithms. Boxes show medians and interquartile ranges. **(e–f)** Cluster quantification errors by StormGraph, ClusterViSu, and DBSCAN relative to ground truth. The fractional error in the percentage of localizations assigned to clusters (e) and the fractional error, Δσ/σ, in the standard deviation, σ, of number of localizations per cluster (f) are plotted *versus* the fractional error, Δμ/μ, in the mean number of localizations per cluster, μ, for each of the 64 simulated ROIs for StormGraph and DBSCAN. For ClusterViSu, the errors are plotted for the 49 ROIs for which analysis was completed in under 2 h. N.b. ±1 means ±100% error, and errors greater than 100% are shown in the shaded regions, which have different axis scales. The colors in (e-f) correspond to the algorithm and parameters used and match the colors in (d): ±uncertainties, *k* = 15 for StormGraph; MinPts = 20, *ε* = 28.2 nm for DBSCAN. These DBSCAN parameters achieved the highest median absolute NMI.

According to NMI and mean F-measure, StormGraph consistently outperformed ClusterViSu and generally performed better than DBSCAN regardless of parameter choice (Figures 2d and S4). DBSCAN’s performance was very sensitive to the choice of parameters and no single choice was suitable for all of the data (Figure S5), demonstrating its unsuitability for analyzing multiple diverse ROIs. In terms of cluster quantification, the errors associated with StormGraph grouped closer to zero than the errors for either ClusterViSu or DBSCAN (Figure 2e–f). For visual appreciation, specific quantification errors and NMI values for the three example simulations in Figure 2a–c are shown in Figure S4. ClusterViSu tended to substantially underestimate the mean and variance of the number of localizations per cluster, presumably because it detected too many small spurious clusters. Quantification errors by DBSCAN were close to zero for many of the ROIs, but the occurrence of some large errors reinforces its unsuitability for analyzing multiple diverse ROIs. Meanwhile, StormGraph was very robust for values of *k* ranging from 10 to 20 when using the kNN method to set *r*_0_, and it was similarly robust for the automated heuristic method of setting *r*_0_ (Figures 2d and S4). Inspection of the quantification errors reveals that cluster quantification by StormGraph was generally more accurate when localization uncertainties were utilized (Figure S4). Even so, NMI and mean F-measure showed that StormGraph outperformed both ClusterViSu and DBSCAN whether localization uncertainties were utilized or not (Figures 2d and S4).

Nonetheless, clustering algorithms cannot be expected to achieve perfect results. Despite the existence of a ground truth in each of our simulated ROIs, three contributions to their realism posed hurdles to algorithms identifying the true clusters. Firstly, distinct ground-truth clusters could appear very close together by chance and therefore be almost indistinguishable as separate clusters. This is apparent in Figure 2a–c. Secondly, molecules were distributed randomly among the nanoclusters, therefore some ground-truth nanoclusters might have molecular densities too sparse to discern as clusters. Thirdly, multiple localization of single molecules, which our simulations included, causes individual molecules to manifest as small clusters of localizations. This increases spurious detection of small clusters among randomly distributed molecules. Although StormGraph tests for and subsequently excludes clusters that cannot be confidently distinguished from multiply localized individual molecules, as described in the Methods, chance instances of two such molecules occurring close together would not be excluded. ClusterViSu and DBSCAN perform no such tests at all, and Figures 2 and S5b show that, for the same minimum cluster size of 5 localizations, StormGraph detects fewer spurious clusters than either. This is not the only reason for StormGraph having the best test results, however, as StormGraph still outperformed ClusterViSu and DBSCAN for simulated data in which every molecule yields exactly one localization (Figure S6). Overall, although clustering results from all algorithms inevitably deviated from ground-truth, StormGraph generally deviated the least.

Notably, StormGraph’s single-level clustering results, to which our reported performance statistics relate, sometimes displayed merging or fragmentation of ambiguous ground-truth clusters. In such instances, clusters closer to ground truth were usually still visually evident in some level of the cluster hierarchy. An example of this is demonstrated in Figure 2c (Levels 1–4). Furthermore, for simulated data with nanoclusters of 50 nm radius, we were able to manually identify a level of clustering from StormGraph’s multi-level output that accurately recovered the ground-truth nanoclusters that composed larger ground-truth aggregations (Figure S7). Thus, StormGraph is able to identify meaningful clusters at multiple scales.

We also compared StormGraph to the Bayesian method of Rubin-Delanchy et al. (29), the only existing algorithm that fully utilizes localization uncertainties. However, we could not test the Bayesian method on the ROIs used in Figure 2, which typically contained between 10^4^ and 10^5^ localizations, due to excessive memory demands. Therefore, we simulated 30 new ROIs of size 1 μm × 1 μm containing fewer than 10^4^ localizations (see Figure 3b for two examples). The Bayesian method requires several user inputs, which are fully described by Rubin-Delanchy et al. We used the default prior distribution of cluster radii because this covered the expected range of possible cluster sizes in our simulations. For the generation of cluster proposals, we used the default range and increment of values of the threshold *T* (5 to 500 in increments of 5) and values from 5 nm to 210 nm in increments of 5 nm for the radius *R.* For the Dirichlet concentration parameter *α* and the prior probability, *p,* of localizations being non-clustered, we started with the default values (20 and 0.5 respectively) and then adjusted them in an attempt to achieve better clustering results. For StormGraph, we again tested both the heuristic method and the kNN method, for which we varied *k* from 10 to 20, to automatically determine *r*_0_. We also tested different values of StormGraph’s per-localization significance parameter *α*. Because the Bayesian method does not require a minimum cluster size, we discarded the MinCluSize parameter from StormGraph, in which case it defaults to 3, the smallest number of points that can physically constitute a “cluster”. Thus, we operated StormGraph with just one or two input parameters.

**Figure 3:**
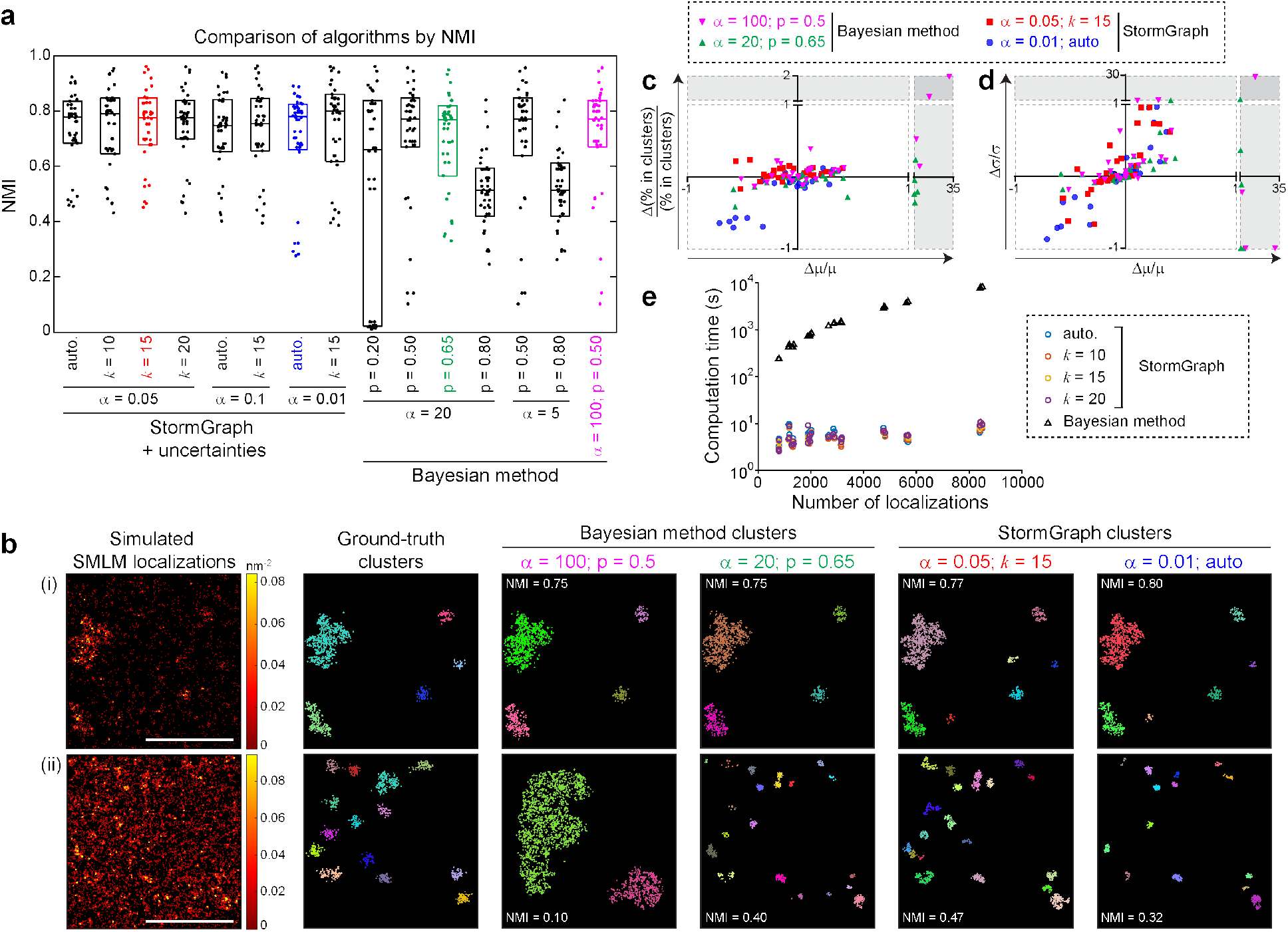
StormGraph is much faster and more suitable for diverse ROIs than the Bayesian method. **(a)** Normalized mutual information (NMI) values measuring the performances of StormGraph and the Bayesian method at assigning localizations to clusters compared to ground truth (NMI = 1 =⟹ 100% match) for 30 simulated 1 μm × 1 μm ROIs. StormGraph was implemented using (+) localization uncertainties and using either the heuristic method (auto.) or the kNN method (*k* = 10, 15, or 20) to set *r*_0_. Three values of the per-localization significance parameter *α* were tested: 0.01, 0.05, and 0.1. For the Bayesian method, the prior *p* and the Dirichlet concentration parameter *α* were varied. See the main text for full details of inputs to the Bayesian method. **(b)** Two examples from the 30 simulated ROIs (left-most panels, color bar = density, scale bar = 500 nm), their ground-truth clusters, and clusters detected by the Bayesian method and StormGraph, each using two different parameter selections. Corresponding NMI values are shown. Top: an ROI with close-to-average NMI values for all clustering results. Bottom: an ROI with relatively poor NMI values for all clustering results. For the Bayesian method, clusters are shown for the tested parameters with the overall best mean performance measured by NMI (magenta) or **best** lowest NMI (green). For StormGraph, clusters are shown for the parameters that we recommend (red) and for the tested parameters with the **worst** lowest NMI (blue). N.B. for the bottom ROI, the figure shows the **worst** StormGraph results (blue parameters) and the **best** Bayesian method results (green parameters). **(c–d)** Cluster quantification errors by StormGraph and the Bayesian method relative to ground truth. The fractional error in the percentage of localizations assigned to clusters (c) and the fractional error in the standard deviation, σ, of number of localizations per cluster (d) are plotted *versus* the fractional error in the mean number of localizations per cluster, μ, for each of the 30 simulated ROIs. N.b. ±1 means ±100% error, and errors greater than 100% are shown in the shaded regions, which have different axis scales. The parameter selections correspond to the highlighted parameters in (a-b). The blue dots in the bottom left of the plots represent the worst StormGraph results achieved by any of the tested parameters, whereas only the best Bayesian method results are shown. **(e)** Computation time taken by StormGraph (+ uncertainties; *α* = 0.05) and the Bayesian method (default parameters but reduced number of cluster proposals) to analyze simulated ROIs *versus* number of localizations. Computations were performed on a standard desktop computer with 16 GB RAM and running Ubuntu 16.04 on a solid-state drive.

By evaluating NMI, mean F-measure, and cluster quantification errors, we found that StormGraph was more robust than the Bayesian method (Figures 3a and S8). Importantly, the Bayesian method parameter values that produced the best results on average also produced extremely poor results for some ROIs that contained greater proportions of non-clustered molecules (Figures 3 and S8). The third column of Figure 3b(ii) shows a clear example of this. We were able to improve the results of the Bayesian method for those ROIs by adjusting parameters, but this decreased the quality of results for other ROIs. None of the tested parameter values enabled the Bayesian method to perform as well as StormGraph overall for the entire set of 30 ROIs. Even for the overall best choices of parameter values, the Bayesian method always yielded substantial quantification errors for some ROIs, whereas the parameter values that we generally recommend for StormGraph (*α* = 0.05, *k* = 15) always produced relatively reliable quantification (Figure 3c–d). The worst results returned by StormGraph occurred when analyzing ROIs with relatively high densities of non-clustered molecules using the heuristic method for setting *r_0_* coupled with a stringent per-localization significance value (*α* = 0.01). This is unsurprising because the heuristic method is not intended for situations with large numbers of non-clustered molecules and a stringent value of *α*, by definition, increases the probability of false negatives. Even so, these settings still gave results that were comparable to the best results of the Bayesian method for most ROIs. Together, our tests indicate that StormGraph is more suitable for analyzing multiple diverse ROIs than the Bayesian method.

Furthermore, we found that StormGraph was 100 to 1,000 times faster than the Bayesian method at clustering each of the 30 simulated 1 μm × 1 μm ROIs (Figure 3e). Extrapolation of the computation times suggests that the Bayesian method could take ~12 hours (ignoring memory constraints) to analyze a single ROI containing ~30,000 localizations on a standard desktop computer, whereas StormGraph could analyze an entire experiment consisting of 30 such ROIs from each of two conditions or cell types (60 ROIs in total) in under 2 hours. For reference, in our lab, a typical dSTORM analysis of BCR clustering involves analyzing one ROI per B cell from > 20 cells per condition, and each ROI usually contains between 10^4^ and 10^5^ (average ~30,000) localizations. Significantly, the long computation time of the Bayesian method prevents adequate tuning of its various parameters. All things considered, StormGraph may be more practical than the Bayesian method for analyzing SMLM data.

Finally, using simulated circular clusters, we compared StormGraph to the H-function, which is derived from Ripley’s *K*-function (22) and often used to summarize clustering in SMLM data (e.g. (2)). Ripley’s *K*(*r*) function measures the average number of points, normalized by global density, within a disk of radius r centered on a point in the data. The L-function square-root normalizes *K*(*r*) so that its expected value is *r* for a uniformly random distribution of points, and *H*(*r*) further subtracts *r* so that its expected value is zero. Significant deviation of *H*(*r*) above zero indicates clustering, and the value of *r* at which *H*(*r*) achieves its peak is often taken to indicate the size of clusters. In our tests, the H-function was biased towards the clusters containing the most points, as is mathematically expected, and, unlike StormGraph, it did not provide an accurate measure of cluster radius (Figure S9).

### StormGraph is robust to changes in global density of SMLM localizations

Because the global density of SMLM localizations can vary between ROIs, batch processing cluster analysis of multiple ROIs, and comparison of results across samples, is only appropriate if the algorithm results are not influenced by the global localization density. This represents a fundamental limitation of the commonly used DBSCAN algorithm, whose user input parameters explicitly define a threshold density that does not adapt to the data. Our tests in Figures 2, S4, S5, and S6 using simulated ROIs spanning a range of localization densities highlighted this and conversely showed that StormGraph is robust to heterogeneity between ROIs. We further showed that StormGraph is robust specifically to global localization density by applying it to a dSTORM ROI, which contained heterogeneous clusters of immunoglobulin M (IgM)-isotype BCRs on the surface of an HBL-1 B cell (see later for details of experiment), after randomly removing 0%, 25%, 50% or 75% of the localizations (Figure 4a). Although small, low-density clusters were eventually lost, the identification and area quantification of large, unambiguous clusters was robust, and the overall distribution of cluster areas was not significantly impacted (*p* > 0.05; Figure 4b).

**Figure 4:**
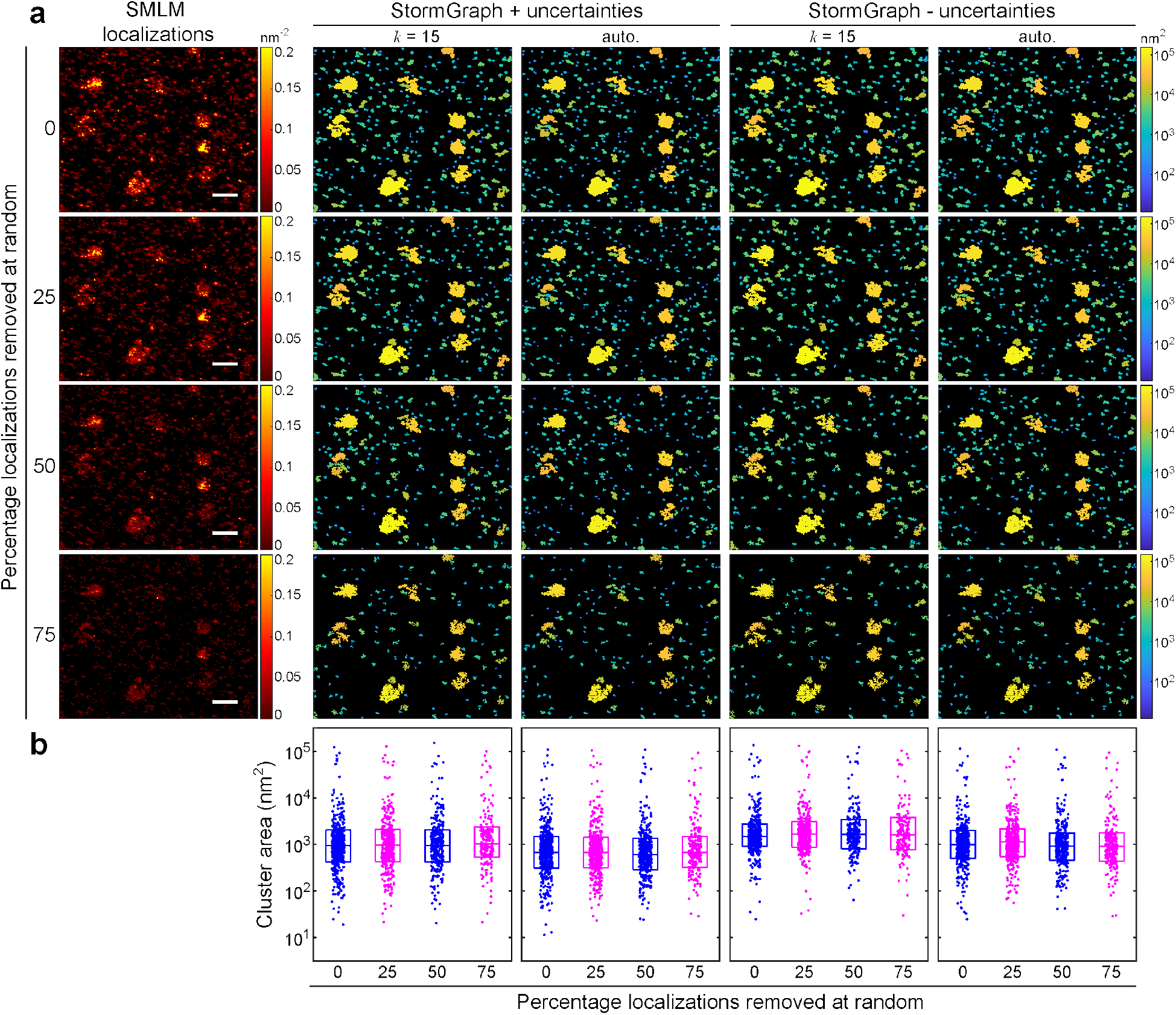
StormGraph results are not sensitive to the global average density of localizations. **(a)** 0%, 25%, 50% or 75% of the localizations were randomly removed from an ROI from real dSTORM data of IgM-BCRs (immunolabeled using Alexa Fluor 647) on an HBL-1 cell (left panels; color bar = density (nm^−2^)). StormGraph was then applied to detect clusters and calculate their areas (remaining panels; color bar = cluster area (nm^2^)). Localization uncertainties were either used (+) or not used (-) during cluster detection and the value of *r*_0_ was set using either the kNN method with *k* = 15 or the heuristic method (auto.). Scale bar = 500 nm. **(b)** Cluster areas quantified by each implementation of StormGraph for each of the four datasets, showing that the distribution of quantified cluster areas was not significantly affected by the random removal of localizations (p > 0.05 as determined by two-sample Kolmogorov-Smirnov tests). Boxes show medians and interquartile ranges.

We also tested StormGraph’s sensitivity to random noise by artificially adding random localizations (with uncertainties) to the unadulterated dSTORM ROI in Figure 4 (Figure S10). StormGraph’s ability to detect all but small, low-density clusters was again robust, and its overall sensitivity to random noise was minimized by including localization uncertainties and using the kNN method to determine *r*_0_. This implementation with *k* =15 resulted in no statistically significant (*p* < 0.05) change in the distribution of cluster areas until the ratio of true localizations to artificial localizations was < 2.

### Quantification of heterogeneous B-cell receptor clustering from 2D dSTORM data using StormGraph

To test StormGraph on real SMLM data, we used it to analyze the clustering of IgM-BCRs on the cell membranes of B lymphocytes. Previous studies found that IgM-BCRs on resting B cells exist in preformed nanoclusters (2; 40). Treatment of resting B cells with anti-Ig antibodies, which are widely used as surrogates for antigens to trigger B-cell activation, alters the spatial arrangement of BCRs, but the exact nature of the alterations remains controversial (2; 40; 41). Formation of larger BCR clusters has been observed upon anti-Ig treatment of resting B cells (2), suggesting a role for increased BCR clustering in the activation of B cells by antigens. Conversely, Reth and colleagues have proposed a model in which the initial step in BCR activation is the dissociation of autoinhibited BCR oligomers, and they observe a decrease in BCR nanocluster size after exposing B cells to antigens or to anti-Ig antibodies (40). Here, we provide new insights revealed by dSTORM and StormGraph, which suggest that the two models might not be mutually exclusive.

Using dSTORM, we imaged fluorescently labeled IgM-BCRs on *ex vivo* murine splenic B cells that were either resting or treated with bivalent antibodies against the BCR’s Igκ light chain. Localization coordinates and their associated uncertainties were computationally determined from the fluorescence data. We then used StormGraph (α = 0.05, MinCluSize = 5 localizations) to batch process the analysis of IgM-BCR clustering in ≥ 24 rectangular ROIs > 1 μm^2^ from separate cells and entirely within cell boundaries (Figure 5a). StormGraph automatically discounted any clusters of localizations that could not be confidently distinguished from overcounted single molecules. Using *k* = 15, StormGraph’s single-level clustering results showed that the mean area of IgM-BCR clusters was significantly larger on anti-Igκ-treated B cells than on resting B cells (Figure 5b(i), *p* < 10^−5^), consistent with the “increased BCR clustering” model of BCR activation. However, we did not observe a uniform increase in the distribution of cluster areas. Instead, the difference was mainly due to a distributional shift towards larger areas of the clusters that were already > 6 × 10^3^ nm^2^, which accounted for approximately the 20% largest clusters in both resting and anti-Ig κ-treated cells (Figure 5b(ii)). The majority of clusters present on anti-Igκ-treated cells were, in fact, small multimers that were comparable to, or even smaller than, the IgM-BCR clusters on resting cells, a prediction of Reth’s dissociation-activation model of BCR activation. The automatic (no *k* value) implementation of StormGraph yielded similar conclusions (Figure S11). Our observations here, powered by StormGraph’s ability to analyze heterogeneous clustering in many ROIs, suggest that the two different models of antigen-induced BCR rearrangement coexist in the same cells. Antigen-induced BCR activation might be associated with both the dissociation of small oligomers and the aggregation of larger nanoclusters.

**Figure 5:**
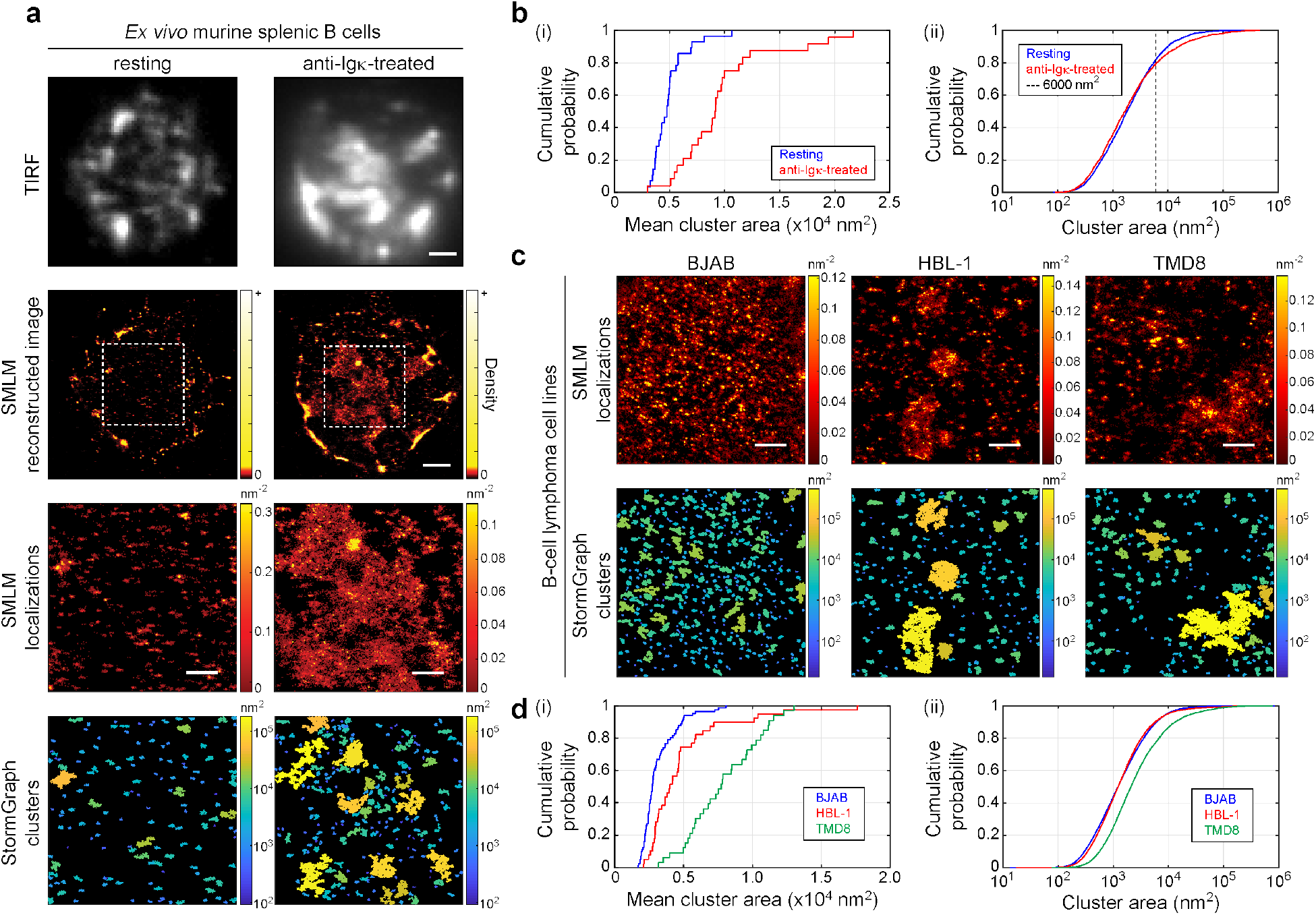
StormGraph analysis of IgM-isotype B-cell antigen receptors (IgM-BCRs) on *ex vivo* murine splenic B cells and human B-lymphoma cell lines imaged using dSTORM. **(a)** IgM-BCRs, immunolabeled using Alexa Fluor 647, on *ex vivo* murine splenic B cells that were either left untreated (resting; left panels) or treated with bivalent anti-Igκ antibodies (anti-Igκ-treated; right panels). Cells were imaged first by total internal reflection fluorescence (TIRF) microscopy (top row) and then by dSTORM (second row; images reconstructed from dSTORM localizations). Scale bar = 1 μm. Third row: IgM-BCR dSTORM localizations in the ROIs (dashed white boxes) in the second row. Scale bar = 500 nm, color bar = localization density (nm^−2^). Bottom row: Clusters identified by StormGraph, colored by their areas (nm^2^). **(b)** Cumulative distribution functions (CDFs) for cluster areas in ROIs from 28 resting (blue) and 24 anti-Igκ-treated (red) *ex vivo* murine splenic B cells. Panel (i) shows the mean cluster area in each ROI (p < 10^−5^). Panel (ii) shows all cluster areas from all ROIs. The increased mean cluster areas in anti-Igκ-treated cells compared to resting cells is due to clusters larger than ~6000 nm^2^. **(c)** StormGraph analysis of IgM-BCRs imaged by dSTORM on resting BJAB, HBL-1 and TMD8 cells. Top: ROIs containing IgM-BCR dSTORM localizations from a representative BJAB Burkitt’s lymphoma cell or from HBL-1 and TMD8 ABC DLBCL cells. Scale bars = 500 nm, color bar = density (nm^−2^). Bottom: Clusters identified by StormGraph, colored by their areas (nm^2^). **(d)** CDFs for cluster areas in ROIs from 81 BJAB (blue), 39 HBL-1 (red), and 33 TMD8 (green) cells. Panel (i) shows the mean cluster area in each ROI (HBL-1 vs BJAB: p < 10^−4^; TMD8 vs BJAB: p < 10^−14^). Panel (ii) shows the areas of all clusters in all ROIs. The larger mean area of clusters on HBL-1 cells compared to BJAB cells is due to small numbers of very large clusters. All StormGraph results shown here were generated using localization uncertainties, *k* = 15, *α* = 0.05, and MinCluSize = 5.

We next used StormGraph to analyze the aberrant spatial arrangement of IgM-BCRs on B-lymphoma cells with an activated B-cell like (ABC) phenotype. Davis et al. showed that chronic BCR signaling is a feature of many ABC-subtype diffuse large B-cell lymphomas (DLBCLs) and, using diffraction-limited microscopy, they observed large IgM-BCR clusters in the absence of any stimulus on the ABC DLBCL cell lines HBL-1 and TMD8 but not on the Burkitt’s lymphoma cell line BJAB (42). To better characterize aberrant IgM-BCR clustering in some ABC DLBCLs, we used dSTORM and StormGraph to assess IgM-BCR cluster areas from ≥ 33 ROIs > 1 μm^2^ on HBL-1, TMD8, and BJAB cells (Figure 5c). ROIs contained between 5 × 10^3^ and 3 × 10^5^ localizations. Using *k* = 15, StormGraph revealed that the mean areas of IgM-BCR clusters on HBL-1 and TMD8 cells were significantly larger than on BJAB cells (*p* < 10^−4^ and *p* < 10^−14^ respectively; Figure 5d(i)). The difference between the distributions of IgM-BCR cluster areas on BJAB and HBL-1 cells resembled the difference between resting and anti-Igκ-treated B cells. Although HBL-1 and BJAB cells both had many small IgM-BCR clusters, the HBL-1 ABC DLBCL cells displayed greater size and/or frequency of large clusters > 10^4^ nm^2^. In contrast, the other ABC DLBCL cell line, TMD8, displayed an overall upward shift in the distribution of cluster areas compared to BJAB, though > 80% of IgM-BCR clusters on the TMD8 cells still had areas < 10^4^ nm^2^ (Figure 5d(ii)). The automatic implementation of StormGraph yielded similar results (Figure S11). Also, although it reduced the magnitude and statistical significance of some of the differences in cluster areas reported by the automatic implementation of StormGraph, ignoring localization uncertainties during StormGraph analysis of our anti-Igκ and B-lymphoma dSTORM experiments did not qualitatively alter results (Figure S11). All together, our observations reveal that IgM-BCRs exist in a heterogeneous combination of small and large clusters in two ABC DLBCL cell lines and that their increased frequencies of large IgM-BCR clusters mimic observations for B cells activated by anti-Ig antibodies. This supports findings by Davis et al. (42) that those ABC DLBCL cell lines exhibit chronic BCR signaling.

### Computation time

To investigate the time complexity of StormGraph, we timed StormGraph clustering for each of the dSTORM ROIs that we analyzed in Figure 5 and plotted the computation times against the total number of localizations per ROI (Figure 6a). Neither the choice of method (kNN or automatic) to determine *r*_0_ nor whether or not localization uncertainties were utilized substantially influenced the computation time. For StormGraph utilizing uncertainties with *k* = 15 and *α* = 0.05, we empirically determined that the computation time, *T*, taken by StormGraph for one ROI containing *N* localizations from our IgM-BCR dSTORM data could be estimated by *T* =1.3 × 10^−4^ × *N*^1.32^ seconds (Figure 6b). The theoretical time complexity of StormGraph is difficult to determine because it depends on many factors, but this empirical relationship indicates a time complexity of approximately 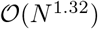. The empirical relationship between *N* and *T* translates to computation times per ROI of < 1 minute for 10^4^ localizations, ~10 minutes for 10^5^ localizations, and ~3 hours for 10^6^ localizations. The cumulative time taken by any of the four tested implementations of StormGraph to analyze all 167 ROIs from our two IgM-BCR dSTORM experiments was under 12 hours. Hence, StormGraph is particularly well suited for analyzing receptor clustering on B and T cells, where an SMLM ROI that occupies a large fraction of the area of a cell would typically contain on the order of 10^4^ or 10^5^ localizations.

**Figure 6:**
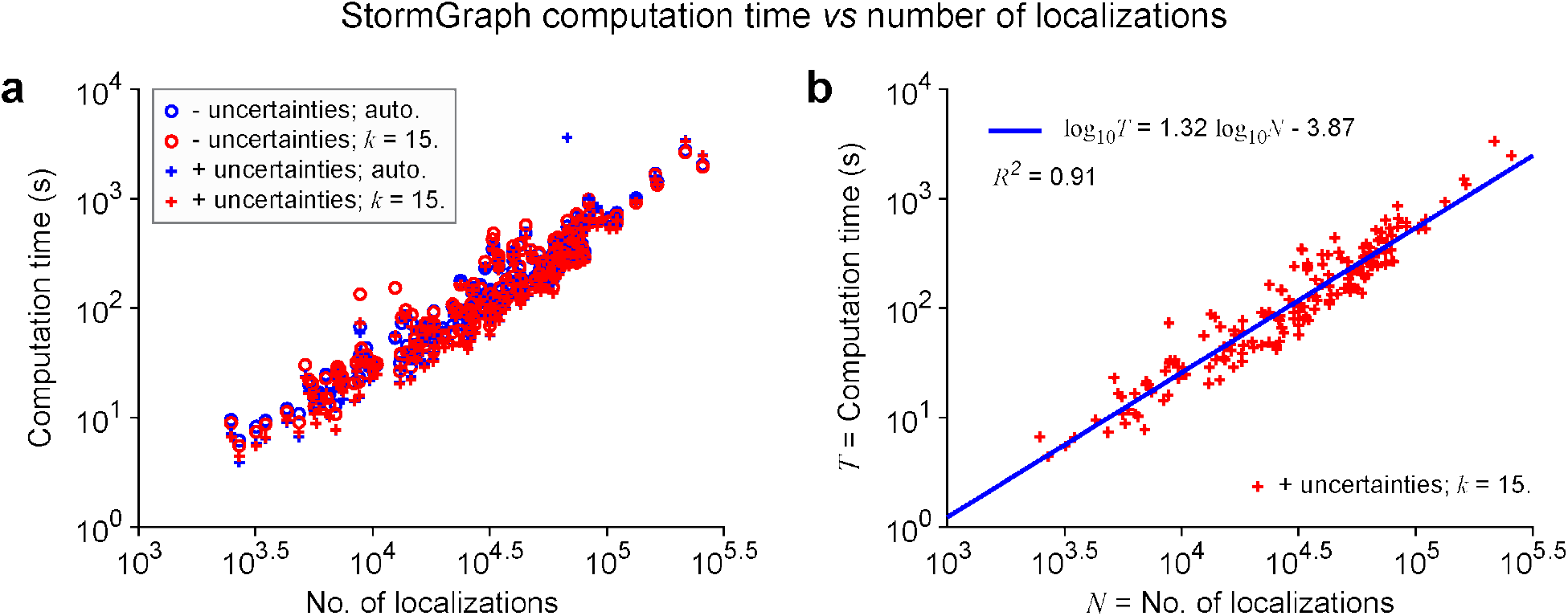
Relationship between computation time for StormGraph and number of localizations in an ROI. **(a)** Scatter plot on logarithmic axes of computation times *versus* number of localization for different implementations of StormGraph. Computations were for the actual dSTORM data analyzed and presented in Figure 5 and were performed on a standard desktop computer with 16 GB of RAM and running Ubuntu 16.04 on a solid-state drive (SSD). **(b)** A linear relationship fitted to the logarithm of computation time *versus* the logarithm of the number of localizations *N* in an ROI shows that StormGraph with our recommended settings (using localization uncertainties, *k* = 15, *α* = 0.05) has an empirical time complexity of approximately 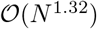.

### Two-color analysis of cluster overlap

Co-aggregation and segregation of different molecules are important cellular mechanisms for regulating signal transduction and can be studied via multi-color SMLM. To quantify colocalization of clusters of molecules labeled by two different colors (e.g. red and blue), the StormGraph software quantifies the total area of overlap divided by each of the following: (1) total red cluster area; (2) total blue cluster area; and (3) total area covered by clusters of either color, yielding the Jaccard index (43) (Figure 7). The software also reports analogous quantities using numbers of localizations instead of areas (not shown). To estimate the maximal experimentally observable colocalization, colocalization analysis should first be applied to the same molecular species labeled with two different probes. This rarely yields 100% colocalization for several reasons, including differing affinities of antibody-fluorophore conjugates, differing photophysical properties of fluorophores, and the inability of two probes to occupy the same binding site.

**Figure 7:**
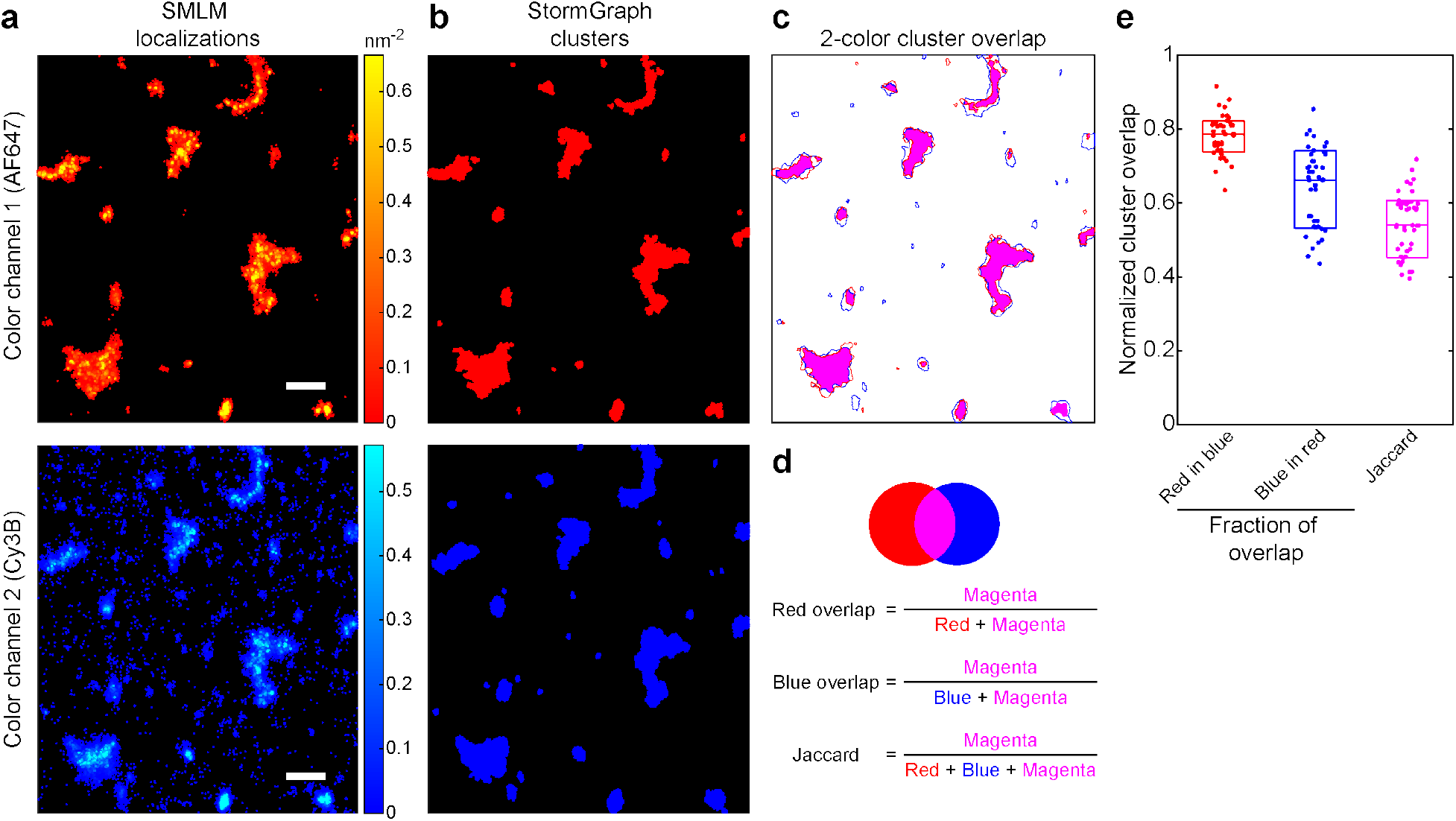
Two-color cluster overlap analysis using StormGraph. **(a)** IgG-isotype B-cell antigen receptors (IgG-BCRs) on A20 B cells were labeled simultaneously with bivalent anti-IgG antibodies that were conjugated to either AF647 (top panel; red) or Cy3B (bottom panel; pseudo-colored blue) and then imaged using dSTORM. Bivalent antibodies were used to induce clustering, since each antibody can bind up to two IgG-BCRs. The IgG-BCR dSTORM localizations in an ROI from one representative cell were analyzed using StormGraph. Scale bar = 500 nm, color bars = density (nm^−2^). **(b)** Binary images of the AF647 (top) and Cy3B (bottom) clusters identified by StormGraph in the ROI shown in (a). **(c)** Merged image of the outlines of the AF647 clusters (red) and Cy3B clusters (blue) identified by StormGraph, with the overlapping areas colored in magenta. **(d)** Pictorial description of the three area-based cluster overlap scores calculated by StormGraph, in the same order as the columns in panel (e). **(e)** Cluster overlap scores calculated using the formulae in panel (d) for 31 StormGraph-analyzed ROIs from multiple A20 cells imaged in the same experiment. Each ROI contributes one dot to each column. Boxes show medians and interquartile ranges. These scores determine the maximum observable overlap that could be expected for clusters of IgG-BCRs and a different molecule labeled using these same two fluorophores on A20 cells, imaged using the same imaging setup and analyzed by StormGraph.

To demonstrate cluster overlap analysis by StormGraph, we performed such a positive control experiment by simultaneously labeling cell-surface IgG-BCRs on murine A20 B cells with anti-IgG antibodies conjugated to either Alexa Fluor 647 (AF647) or Cy3B fluorophores. These antibodies were bivalent, thus inducing formation of large clusters prior to cell fixation. Both color channels were imaged using dSTORM and aligned using custom MATLAB code to correct for chromatic aberrations. We then analyzed multiple ROIs using StormGraph (Figure 7). On average, we found 79% overlap of the IgG-AF647 clusters with the IgG-Cy3B clusters and 66% overlap of the IgG-Cy3B clusters with the IgG-AF647 clusters (Figure 7e). This difference is likely due to differing qualities of the AF647- and Cy3B-conjugated antibodies. The Jaccard index cannot exceed either one-sided overlap score, and we obtained an average Jaccard index of 0.5. In a similar experiment staining tubulin, Andronov et al. obtained ~40% overlap of each probe with the other using ClusterViSu (25). This shows that StormGraph performs well as part of a pipeline for analyzing cluster colocalization by SMLM.

### Clustering in three dimensions

To extend a clustering algorithm to 3D SMLM data, two challenges must be addressed. First, most 3D SMLM techniques achieve lower axial resolution than lateral resolution. However, StormGraph implicitly assumes that all dimensions should be weighted equally during graph construction. Therefore, StormGraph pre-processes 3D data for cluster identification, but not subsequent quantification, by rescaling the axial (z) dimension so that average axial and lateral positional uncertainties, when known, become equal. Second, 3D SMLM localizations are often concentrated around a focal plane, causing their axial distribution to be nonuniform. To account for this, StormGraph now uses the parameter *α* to obtain a *z*-dependent node-degree threshold from random point clouds with normally distributed *z*-coordinates (Methods). For situations with localizations distributed uniformly in *z*, StormGraph still retains the option to use a constant threshold instead. On the other hand, DBSCAN is unable to adapt to axial variation in localization density.

We compared the performances of StormGraph and DBSCAN in 3D (“StormGraph-3D” and “DBSCAN-3D”) using simulated 3D data. As in 2D, we found that, overall, the output clusters were closer to ground truth for StormGraph than DBSCAN regardless of parameter choices (Figure S12). We also performed 2D clustering (“StormGraph-2D” and “DBSCAN-2D”) of the *xy*-projections of our simulated 3D data. Including the *z*-component of 3D data generally improves clustering accuracy because localizations and clusters that are separated only in z are inseparable in the *xy*-projection. StormGraph-3D produced the best overall clustering results, but even StormGraph-2D produced better results than both DBSCAN-3D and DBSCAN-2D. Moreover, these results were obtained using the same parameter values for both StormGraph3D and StormGraph-2D, whereas DBSCAN-3D and DBSCAN-2D necessitated different parameter values, as expected. Hence, it is easy to switch between 2D and 3D analyses with StormGraph.

To illustrate StormGraph’s application to real 3D SMLM data, we used dSTORM to image intracellular lysosomal-associated membrane protein 1 (LAMP-1). We simultaneously immunostained LAMP-1 in B16 melanoma cells with two different labels, AF647 and Cy3B, and applied StormGraph (*k* = 15, *α* = 0.1, MinCluSize = 5 localizations) to a 3D ROI with axial variation in localization density and known localization uncertainties (Figure 8a–b). StormGraph detected 363 LAMP-1 AF647 clusters and 129 LAMP-1 Cy3B clusters (Figure 8c–d). The AF647 clusters had volumes ranging from 1.5 × 10^3^ nm^3^ to 7.1 × 10^7^ nm^3^ with a median of 3.5 × 10^5^ nm^3^, and Cy3B clusters had volumes ranging from 3.1 × 10^3^ nm^3^ to 3.7 × 10^7^ nm^3^ with a median of 9.0 × 10^5^ nm^3^ (Figure 8e). The discrepancy in cluster volumes was likely caused by variance in labeling or probe detection. Indeed, we detected almost four times as many AF647 localizations as Cy3B localizations (9.0 × 10^4^ versus 2.5 × 10^4^) and it is evident from Figure 8a–b that Cy3B was inadequate for some of the features observed using AF647. This shows that probe choice is an important consideration for SMLM. Nevertheless, StormGraph detected similar larger-volume clusters in both color channels provided that they were sufficiently labeled by both probes.

**Figure 8:**
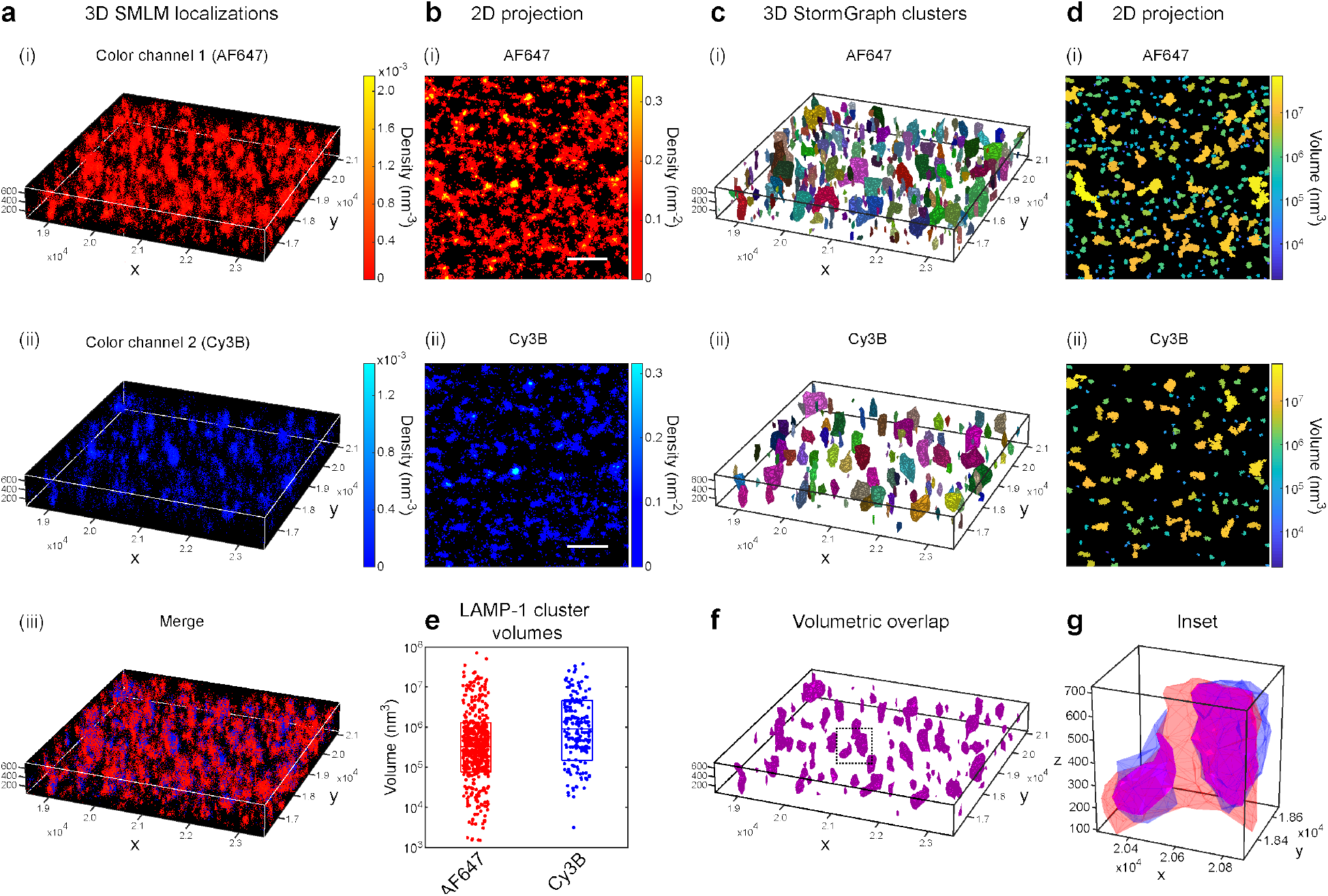
Analysis of 3D SMLM data using StormGraph. **(a)** Localizations of intracellular LAMP-1 labeled simultaneously by two different fluorophores in a B16 murine melanoma cell imaged by two-color 3D dSTORM. LAMP-1 molecules were stained using anti-LAMP-1 primary antibodies and a 1:1 mixture of the same secondary antibody conjugated to either (i) AF647 (red) or (ii) Cy3B (pseudo-colored blue). A 5 μm × 5 μm × 700 nm ROI from one cell was selected for analysis. Color bars = density (nm^−3^). (iii) Merge. **(b)** 2D projections of the (i) AF647 and (ii) Cy3B localization data onto the *xy*-plane. Color bars = density (nm^−2^). Scale bar = 1 μm. **(c)** (i) AF647 and (ii) Cy3B clusters found by StormGraph using localization uncertainties, *k* =15, *α* = 0.1, and MinCluSize = 5 localizations. Clusters of localizations that could not be confidently distinguished from a single, multiply counted fluorescent probe were automatically removed by StormGraph. Colors distinguish different clusters. **(d)** 2D projections of the clusters shown in panel (c) colored according to their 3D volumes (nm^3^). **(e)** All volumes, as colored in (d), of individual AF647 and Cy3B clusters detected by StormGraph. Boxes show medians and interquartile ranges. **(f)** Overlapping volumes (magenta) of the AF647 and Cy3B clusters in panel (c). **(g)** Enlarged region showing overlap (magenta) between one AF647 cluster (red) and two Cy3B clusters (pseudo-colored blue).

Additionally, we computed volumetric 3D overlap between AF647 clusters and Cy3B clusters (Figure 8f–g). Though only 31% of the total AF647 cluster volume overlapped with Cy3B clusters, we found that 50% of the total Cy3B cluster volume overlapped with AF647 clusters. The Jaccard index (which cannot exceed the smaller overlap score, i.e. 0.31) was 0.24. These observations were probably explained by weaker labeling or detection with Cy3B than AF647 such that some clusters lacked Cy3B signal. To our knowledge, our software is the first to compute volumetric overlap for two-color, 3D SMLM data. StormGraph thus offers an alternative to Coloc-Tesseler (44), an extension of SR-Tesseler, for analysis of co-clustering of molecules. Our results for LAMP-1 show that StormGraph can identify and quantify clusters of localizations in 3D SMLM ROIs and, furthermore, that it can detect and quantify overlap between 3D clusters in two-color data.

## Discussion

By converting 2D or 3D SMLM data into a neighborhood graph, StormGraph leverages concepts from graph theory, especially community detection, to assign localizations to clusters at multiple scales. StormGraph can be downloaded from [URL to be inserted upon acceptance for publication] and is run in MATLAB using either a script or a simple graphical user interface (GUI). The software automatically quantifies clusters at an automatically selected scale, quantifies cluster overlap for two-colour SMLM data, and includes MATLAB functions for data visualization in 2D or 3D. Through extensive testing using simulated data, we showed that StormGraph is generally more accurate than existing cluster analysis methods. Furthermore, unlike the popular DBSCAN (23) and methods based on Voronoi diagrams (24; 25), StormGraph can utilize the uncertainties in individual localization positions to enhance clustering accuracy. Previously, only a Bayesian method (29; 30) had this capability, but StormGraph requires fewer user inputs, makes fewer assumptions, and is between 100 and 1,000 times faster. As summarized in Table S1, StormGraph combines several features that are not all available simultaneously in any other SMLM cluster analysis method.

A crucial feature of StormGraph is its automatic determination of scale-dependent thresholds from scale-independent input parameters, whose selection we provide guidelines for in the Methods. Firstly, StormGraph determines a neighborhood radius either based on a user-specified number of neighbors, k, that neighborhoods of clustered localizations should exceed (recommended method) or heuristically without user input (suitable for data in which at least two thirds of the localizations clearly belong to clusters). Secondly, StormGraph decides which localizations are sufficiently dense to be clustered via a user-specified per-localization significance level *α*. Hence, StormGraph’s input parameters — *k* (optional) and *α* (required) — do not require any *a priori* knowledge about quantities that often vary between ROIs in SMLM experiments, such as the localization density, which affects DBSCAN, or the fraction of localizations that do not belong to clusters, which affects the Bayesian method. Consequently and importantly, StormGraph enables unbiased analysis of disparate datasets using identical parameter values, whereas DBSCAN and the Bayesian method do not. Therefore, StormGraph is more suitable than either DBSCAN or the Bayesian method for analyzing SMLM datasets consisting of multiple ROIs and for making comparisons between different conditions, molecules, or cell types.

By applying StormGraph to actual dSTORM data, we simultaneously detected both very small and very large IgM-BCR clusters on the cell membranes of activated B cells. Two opposing models of antigen-induced BCR activation have been proposed by others: one model involves increased aggregation of BCRs and the other involves a decrease in BCR clustering via dissociation. Our observations, quantified by StormGraph, permit a new hypothesis that both models occur together. Perhaps large clusters form from BCRs that must first dissociate from pre-existing small clusters. This would unite the two opposing models for the first time and warrants further investigation. By providing improved quantitative characterization of heterogeneous receptor clustering, StormGraph should enable new insights into the relationship between receptor clustering and receptor signaling.

Note that we deliberately avoided making any statements about numbers of molecules during our analysis of experimental dSTORM data. In SMLM, the ratio of localizations to molecules is not one-to-one. Repeated blinking of fluorophores and labeling of individual molecules by multiple fluorophores cause overcounting of molecules. Spatially unresolvable fluorophores that blink simultaneously cannot be localized and therefore cause undercounting of molecules, which is especially an issue for dense clusters. Undercounting also arises from other experimental sources of error, such as incomplete labeling of molecules or incomplete detection of fluorophores. For well controlled PALM experiments with minimal undercounting, the number of molecules can be estimated (45; 46), but for general SMLM experiments, especially dSTORM using fluorescent immunolabeling of molecules as we performed, accurate determination of the number of molecules per cluster remains a challenge. Therefore, to avoid false biological interpretation of the data, we chose to report only cluster areas or volumes.

During the development of StormGraph, Khater et al. also presented a method to analyze SMLM data using graphs (31). We note important differences between their approach and StormGraph. Both methods filter out non-clustered data points using a node-degree threshold obtained from random data, but whereas StormGraph obtains this threshold using a per-localization significance level *α*, Khater et al. use a parameter equivalent to the *α* parameter in SR-Tesseler. For clustering, StormGraph uses graph-based community detection whereas Khater et al. use the mean shift algorithm (47), an unrelated density estimation method, which is sensitive to a difficult-to-select user-specified bandwidth parameter (48). Khater et al. then use multi-threshold network analysis to identify modular structures within the clusters, whereas StormGraph identifies clusters and their multiple levels of constituent subclusters simultaneously using automated multi-level clustering. Finally, unlike StormGraph, the method of Khater et al. does not account for uncertainties in localization positions.

It should be noted that the uncertainties in localization positions, which result from the finite resolution of SMLM, cause the localization clusters to be larger than the true underlying molecular clusters. StormGraph does not correct for this, nor do any of the other clustering algorithms. Users should therefore be aware that the cluster areas or volumes reported by StormGraph will be slight overestimates of the actual sizes of the molecular clusters. The quantified overlap of localization clusters will also differ slightly from the true overlap of molecular clusters. If the clusters output by StormGraph happen to be approximately Gaussian, then mathematical correction methods (49) could be applied in order to improve estimates of cluster size and overlap. Without any such correction, it is important to perform all imaging at the same resolution in order to keep errors consistent and enable fair comparisons. Nonetheless, StormGraph will advance cluster analysis in the SMLM field. Parameter selection is simple (we recommend *k* = 15 and values of *α* between 0.01 and 0.1 for most data), and a simple MATLAB GUI and script make StormGraph accessible to a wide range of users. With its unique combination of features — including utilization of localization uncertainties and generation of nested clusters across multiple scales — and greater robustness and accuracy than existing algorithms, we believe that StormGraph provides a one-stop shop for SMLM cluster analysis.

## Methods

Calculation of the length scale *r*_0_

### (1) The fully automatic, heuristic method

To automatically determine a length scale *r*_0_ without user input, we implement a variation of the elbow method heuristic. For values of *ε* ranging from 0 to a sufficiently large value based on the optimal affinity scale stated by Arias-Castro (50), we construct the *ε*-neighborhood graph for the data. We then plot the number of connected components (including singletons) against *ε*. This must be monotonically decreasing and typically bears resemblance to a decaying exponential or logistic function. As *ε* increases, an “elbow” region occurs as rapid linking of nodes within clusters at small values of *ε* transitions to slower linking of distinct clusters and dispersed nodes at larger values of *ε*. Eventually all nodes would belong to a single connected component.

Sometimes, a natural number of clusters will be evident as a horizontal (i.e. constant) plateau occurring at > 1 connected component in this plot. In such cases, we find the plateau corresponding to the largest fold increase in the area or volume of the *ε*-neighborhood. Let *ε*_1_ be the value of *ε* at the start of this plateau, and let *ε*_2_ = 2^1/*d*^*ε*_1_, where *d* is the dimensionality of the data, be chosen such that the *ε*_2_-neighborhood is twice the area or volume of the *ε*_1_-neighborhood. If the *ε*_1_- and *ε*_2_-neighborhood graphs have the same number of connected components, then we set *r*_0_ = *ε*_2_ (Figure S1).

Otherwise, we fit a curve *f*(*ε*) to the number of connected components versus *ε* (Figure S1). We choose *f* (*ε*) to be the sum of a constant *b* and either one or two generalized logistic functions of the form

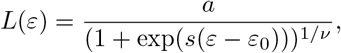

where *b* ≥ 0, *a* ≥ 0, *s* ≥ 0, *ν* > 0, and *ε*_0_ are coefficients to be fit. To avoid overfitting, we only include the second logistic function if it yields a substantial improvement in the goodness of fit and we restrict its allowable values of *ν*. The elbow of this curve is not mathematically well defined, but intuitively it is related to the concavity: the curve achieves maximum (positive) concavity as it approaches the elbow region, and then its concavity decreases as it traverses the elbow region. StormGraph chooses the length scale *r*_0_ to be towards the end of the elbow region as follows. Let *ε*_max_ be the value of *ε* at which *f*”(*ε*), the concavity of *f*(*ε*), is maximized. StormGraph sets *r*_0_ to be the value of *ε* > *ε*_max_ where *f*“(*ε*) first falls below 2% of its maximum value (Figure S1). We chose this 2% threshold empirically after experimenting with different values. Since StormGraph generates a multi-level clustering, and since we have developed and implemented a method to return a single-level clustering based on the hierarchy of clusters, it is safer to overestimate than underestimate a suitable value for *r*_0_. Erroneous merging of clusters can generally be resolved by moving to a finer level of the cluster hierarchy, but erroneous failure to merge clusters cannot be retroactively fixed using the hierarchy. Through experimentation, we found that a 2% threshold worked well while being a generally safe choice.

When localization uncertainties are available in the data, they are initially excluded when utilizing the elbow method to set the initial length scale *r*_o_, which is used for classifying localizations as either clustered or unclustered. The uncertainties are subsequently taken into account during the final use of the elbow method, which sets the value of *r*_o_ that is used for construction of the final graph following elimination of unclustered localizations. Specifically, the graph in which we count the number of connected components for a given *ε* is constructed from Monte Carlo simulated realizations of the data with two nodes connected to each other by an edge if and only if they are within a distance *ε* of each other in at least 75% of the Monte Carlo simulations (see later in the Methods for details of Monte Carlo simulations and edge pruning). Note that edge weights are not relevant here because they do not affect the number of connected components.

### (2) The kNN method

To determine the length scale *r*_0_ for a selected ROI using a *k*-nearest neighbors (kNN) approach, StormGraph first finds the distance of every point in the ROI to its *k*^th^ nearest neighbor. If localization uncertainties are available in the data, this is performed for 100 Monte Carlo simulated realizations of the data, and the 95% confidence level for the *k*^th^ nearest neighbor distance is obtained for every localization. The distribution of *k*^th^ nearest neighbor distances is also obtained for Monte Carlo simulations of random data with the same global average point density as the ROI. A histogram of *k*^th^ nearest neighbor distances should initially increase more rapidly for clustered data than for random data, but the histograms for clustered and random data will eventually intersect each other (Figure S1). Inspired by the automated version of ClusterViSu (25), StormGraph defines *r*_0_ as the distance at which these histograms of *k*^th^ nearest neighbor distances first intersect. Points closer than *r*_0_ to their *k*^th^ nearest neighbor are more likely to exist in clustered data, while points farther than *r*_0_ from their *k*^th^ nearest neighbor are more likely to exist in random data. Moreover, points in clusters will tend to have more than *k* neighbors within a distance *r*_0_, while randomly distributed points will tend to have fewer than *k* neighbors within a distance *r*_0_. However, if this first histogram intersection occurs after the median of the random data’s histogram, this indicates that, on average, the real data is actually more dispersed than the random data, and in this case StormGraph defines *r*_0_ simply as the median of the random data’s *k*^th^ nearest neighbor distances.

### Simulating multiple data realizations and calculation of graph edge weights

StormGraph uses Monte Carlo simulations to simulate multiple realizations of the data by resampling each localization’s coordinates. The new *x, y* and, if applicable, *z* coordinates for a particular localization are drawn independently from normal distributions centered at the original observed localization position. The standard deviations are equal to the corresponding uncertainties recorded in the data. StormGraph then determines the graph edge weights *W_ij_* = 〈*s_ij_*〉 from the Monte Carlo simulations by calculating 〈*s_ij_*〉 to be the mean of the simulated values of *s_ij_* for each specific node pair {*i, j*}.

### Thresholding of node degrees to eliminate unclustered nodes

Setting *α* = 1 skips the thresholding step altogether, allowing all nodes to be considered for clustering. Otherwise, to set the node-degree threshold, StormGraph first constructs *r*_0_-neighborhood graphs with edge weights *s_ij_* for simulated random point clouds with the same global average point density as the SMLM data. For 2D data (and for 3D data with uniform axial acquisition), the random points are uniformly distributed in *x* and *y* (and *z*). Then StormGraph sets the degree threshold as the ((1 – *α*) × 100)^th^ percentile of the aggregated degree distribution of the random simulations. For 3D data with localizations concentrated around a focal plane, StormGraph simulates random data with *z*-coordinates that are distributed normally with the same interquartile range as the data. StormGraph then obtains a *z*-dependent node-degree threshold by fitting a Gaussian curve to node degree versus *z* for the simulated random points and finding the (1 – *α*) × 100% confidence upper bound curve. Thus, for both 2D and 3D data, an expected *α* × 100% of nodes in any of the random simulations would have degrees exceeding the threshold.

For actual data, because the edge weights are calculated by averaging *s_ij_* over Monte Carlo simulations, the number of localizations that would be classified as clustered in random data would usually be less than *α* × 100%. Hence, this averaging using localization uncertainties reduces the detection of spurious, small clusters arising from random spatial fluctuations in density.

If localization uncertainties are not known, then we take a different approach to reduce detection of spurious clusters. Preliminary clusters are defined using a community detection algorithm. A node is then classified as unclustered if it meets any of the following four criteria: (1) it belongs to a preliminary cluster whose mean degree is below the threshold; (2) its own degree is below the threshold and is also a lower outlier (< lower quartile (LQ) −1.5 × interquartile range (IQR)) for its preliminary cluster; (3) its own degree passes the threshold but is a strong lower outlier (< LQ – 3 × IQR) for its preliminary cluster; (4) its own degree is less than half of the threshold. The first criterion provides robustness by spatially averaging node degrees over small areas. This prevents the inclusion of spurious, small clusters. The other three criteria prevent the inclusion of nodes that are visually separate from a cluster but still within a distance *r*_0_ of one.

To avoid biases arising from the choice of algorithm used for the preliminary clustering, StormGraph performs this twice, independently, using two different community detection algorithms, and it then classifies nodes as unclustered if either method does. The two algorithms used are the two-level version of Infomap (51) and the Louvain method (52), which are two of the top performing community detection algorithms (33). Infomap is an information theoretic algorithm based on flow on the graph, while the Louvain method is one of several algorithms that aims to maximize a property of the graph called “modularity”. See Supplementary Note 1 for further technical details.

### Edge pruning

When localization uncertainties are used in the StormGraph algorithm, we prune edges from the final graph that is constructed from only the nodes that are retained after thresholding node degrees. To do this, we delete every edge that has nonzero *s_ij_* in fewer than 75% of the Monte Carlo simulations that were used to calculate the edge weights. This guarantees that any pair of retained edges have at least an estimated 50% probability of co-occurring in the *r*_0_-neighborhood graph for any realization of the data, and the unknown true localization positions is one possible realization. This prevents the linking of clusters that are disconnected in most realizations of the *r*_0_-neighborhood graph but connected in the average graph, since linking of two clusters requires at least one node to be connected by edges to nodes in both clusters simultaneously. After pruning, two clusters can only be linked if they are connected in at least half of the Monte Carlo simulations.

### Merging clusters at the top of the multi-level Infomap hierarchy

To facilitate the identification and quantification of particularly large clusters, StormGraph creates an additional level at the top of the multi-level Infomap cluster hierarchy, if possible, by merging sufficiently interconnected clusters. It is natural to consider the connected components of a graph to be the clusters at the coarsest level of a cluster hierarchy. We therefore use this concept to define the top level of StormGraph’s cluster hierarchy by merging Infomap clusters that form connected components. However, due to the uncertainties in SMLM data, StormGraph only merges clusters if they form stable connected components, which we define as connected components that would remain connected following the random removal or displacement of any one node. Oftentimes, this step results in no merging of clusters and so no additional level of clustering is created.

### Algorithm to obtain single-level clustering from cluster hierarchy

Although various methods exist to select one level from a cluster hierarchy, for example silhouette scores (53) and the gap statistic (54), existing methods are either very computationally intensive or otherwise incompatible with StormGraph. We therefore developed our own fast algorithm to obtain a single-level clustering from the cluster hierarchy output by StormGraph, which we describe here.

The multi-level clustering output by StormGraph is generated from an *r*_0_-neighborhood graph. An alternative type of graph commonly used for clustering problems is the symmetric *k*-nearest neighbor (kNN) graph, in which two nodes are connected by an edge if either of them is among the *k* nearest neighbors of the other. A related graph is the mutual kNN graph, a subgraph of the symmetric kNN graph, in which two nodes are connected by an edge if and only if each node is among the *k* nearest neighbors of the other. One simple clustering algorithm would be to identify the connected components in a symmetric kNN graph or in a mutual kNN graph, where *k* is an adjustable parameter.

In a symmetric kNN graph, it is guaranteed that every node has at least *k* edges. However, as *k* increases, nodes in low-density regions between two distinct clusters quickly become connected to both clusters, while the high-density regions inside the clusters may remain fragmented into multiple connected components until higher values of *k*. A mutual kNN graph, in which every node is guaranteed to have at most *k* edges, more faithfully represents such clusters by preventing nodes in low-density regions from making too many connections. However, mutual kNN graphs often suffer from having singletons and small connected components due to the weak connectivity. We therefore chose to combine the concepts of both the symmetric kNN and mutual kNN graphs.

For a set of points *V* and positive integers *M* and *K* > *M*, we define *G_M,K_*(*V*) to be the union of the symmetric *M*NN graph and the mutual *K*NN graph for vertices *V*. This is still a subgraph of the symmetric *K*NN graph, but it has stronger connectivity than the mutual KNN graph by guaranteeing that every node has at least *M* edges, which in turn ensures that *G_M,K_*(*V*) contains no connected components with fewer than (*M* + 1) nodes.

For each cluster at the top level of the cluster hierarchy, StormGraph decides whether to split the cluster into its subclusters at the next level down in the hierarchy according to the algorithm described below. If the split is rejected, then StormGraph keeps the current cluster and does not examine any of the finer levels of the hierarchy within that cluster. If the split is accepted, then this process is repeated recursively for each of the newly accepted subclusters. A split is automatically rejected if more than 1% of the points in the cluster belong to subclusters with fewer than the minimum number of points, specified by the user, that constitute a cluster.

Let *V* be the set of nodes in a cluster *C*, let 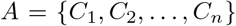 be the set of *n* subclusters of *C* at the next finest level of the cluster hierarchy, and let 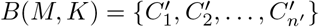 be the set of *n*’ connected components of the graph *G_M,K_*(*V*). StormGraph decides whether to split cluster C into its constituent subclusters A using the following algorithm:

1. Construct *G*_2,*K*_(*V*) for all integers *K* ∈ {6, …,*K*_1_}, where *K*_1_ is the smallest integer such that *G*_2,*K*1_(*V*) is connected. We empirically chose the minimum value of *K* to be 6 because this usually results in randomly distributed points forming a single connected component.
2. Find the value of *K* for which *B*(2, *K*) is most similar to *A* according to some measure of similarity. Denote this value of *K* by *K*\*.
3. Split cluster *C* into subclusters *A* if the similarity between *A* and *B*(2, *K*\*) is greater than both a threshold similarity and the similarity between *C* and *B*(2, *K*\*).

The most obvious choices for a similarity measure to score the similarity between two clusterings of the nodes V are normalized mutual information (NMI) (38) and mean F-measure (39). We require a similarity measure that is defined even if one of the clusterings being compared consists of only a single cluster. This eliminates NMI as a suitable choice, so we use mean F-measure.

Let *F*(*A, B*) denote the similarity of clustering *A* to clustering *B* as measured by the mean F-measure. The F-measure or *F*_1_ score for a binary classification problem in which a cluster *C_i_* is compared to a reference cluster 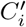 (usually the ground-truth cluster that the cluster *_i_*, found by a clustering algorithm, is supposed to recover) is defined as the harmonic mean of precision (*P*) and recall (*R*):

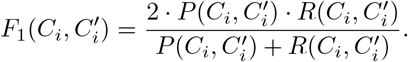

The precision 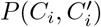 is the fraction of *C_i_* that belongs to 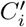, and the recall 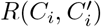 is the fraction of 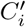 that belongs to *C_i_*. The mean F-measure *F*(*A, B*) is then defined as the weighted arithmetic mean of the maximum F-measures for each of the clusters 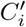 in *B*:

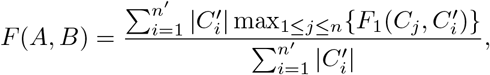

where 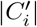 denotes the number of points in 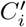.

The mean F-measure is not symmetric, i.e. *F*(*A, B*) ≠ *F*(*B,A*), which is not desirable in our situation where we wish to compare two clusterings, neither of which is necessarily ground-truth. To avoid having to choose one of the clusterings *A* and *B* to be the reference, we define a symmetric similarity measure, 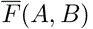, as the arithmetic mean of *F*(*A, B*) and *F*(*B, A*):

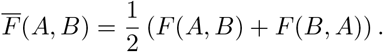

This is the similarity measure that we use in our algorithm for obtaining a single-level clustering from the hierarchy. It ranges from 0 to 1, and 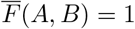 if and only if *A* and *B* are identical. We impose a minimum similarity score of 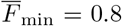 for a cluster split to be considered. Thus, we split cluster *C* into its highest level of subclusters, *A*, if *A* is at least 80% similar to *B*(*M, K*\*) and is also a closer match to *B*(*M, K*\*) than the single, unified cluster *C* is. The 80% similarity threshold prevents the fragmentation of a cluster if there is not substantial consensus between the two independent subclusterings. This threshold could be tuned to make it more or less difficult to split a cluster into finer levels of subclusters. In particular, a threshold of 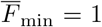 would demand perfect agreement between the subclusters of *C* and the alternative, independent clustering *B*(*M, K*\*) for the subclusters to be accepted as a better clustering of *V* than a single cluster. We chose a threshold of 0.8 to allow some leniency.

### Identifying clusters that can be confidently distinguished from multiply counted single molecules

Localizations arising from multiply counted single molecules may be falsely identified as clusters. As an optional step during StormGraph analysis, clusters of localizations that cannot be distinguished with high confidence, due to their positional uncertainties, from multiply counted single molecules can be identified and subsequently reclassified as unclustered (cluster label 0). To do this, StormGraph checks each cluster systematically as follows.

First, for each pair of localizations, **X**^i^ = (*x^i^, y^i^, z^i^*) and **X**^j^ = (*x^j^, y^j^, z^j^*), in the cluster, let **Y**^ij^ = **X**^i^ – **X**^j^ be their vector difference, and let **Σ**^ij^ be the covariance matrix for the coordinates of **Y**^ij^. The off-diagonal elements of **Σ**^ij^ are assumed to all be 0 (i.e. the uncertainty in each coordinate of a localization is assumed to be independent of its other coordinates). Assuming each molecule to be approximated by a point particle of zero size, the *m*^th^ diagonal element of **Σ**^ij^ is 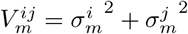, where 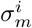 denotes the standard deviation for the uncertainty in the m^th^ coordinate of localization i, as given by the input data.

This assumes that the true position is identical for all localizations originating from the same molecule. In practice, the fluorophore positions may be different from the actual molecule positions. For example, when molecules are detected using antibodies, the fluorophore conjugated to the antibody may be located as much as 10 nm away from the antibody’s binding site. In addition, if each molecule can be labeled by more than one fluorophore, then the true positions of localizations originating from a single molecule will not only be different from the actual molecule but also from each other. If the sizes of the molecule and fluorescent label are not negligible, they can be approximately taken into account in the following way. For mathematical simplicity, we approximate the uncertainty due to the molecule and label size as an isotropic Gaussian distribution with variance (*r*/3)^2^, where *r* is the effective radius of the molecule and fluorescent label combined, which is specified by the user based on underlying biophysical knowledge. We then add this variance term twice (once each for localizations **X**^i^ and **X**^j^) to each of the diagonal elements in **Σ**^ij^. For our simulated data, this was not necessary as the true position of every localization was at the centre of a simulated molecule. For our BCR dSTORM data, we used *r* = 8 nm.

Next, we construct the statistic 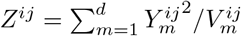 for each pair of localizations, where d is the number of dimensions (2 or 3) and 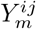 denotes the *m*^th^ coordinate of the vector **Y**^ij^. If two localizations **X**^i^ and **X**^j^ have the same true position, then *Z^ij^* is chi-squared distributed with *d* degrees of freedom. We then look for pairs of localizations for which *Z^ij^* exceeds a desired quantile of the appropriate chi-squared distribution, indicating confidence that they originated from different molecules. Because we are testing multiple pairs of localizations for significance, we correct for multiple hypothesis testing using the Šidák correction. If we desire a significance level of 1 – *q*, then we look for pairs of localizations for which *Z_ij_* exceeds the (*q*^1/*N*^)^th^ quantile of the chi-squared distribution with *d* degrees of freedom. Here, *N* is the number of localizations in the cluster. Even though there are *N*(*N* – 1)/2 pairs of localizations, the null hypotheses are that each localization originated from the same molecule as all other localizations in the cluster, and so there are only *N* hypotheses to test. By default, StormGraph uses a significance level of 0.05, so it uses the (0.95^1/*N*^)^th^ quantile. Finally, since a cluster must always contain at least three localizations (we do not consider pairs of localizations to be clusters), StormGraph increases confidence further by demanding that at least two localizations are each, probabilistically, sufficiently far from at least two other localizations. This way, a single outlying localization within a cluster is not sufficient on its own to qualify the cluster as containing multiple molecules with high confidence.

### Guidelines for StormGraph parameter selection

StormGraph has three user-controllable parameters. The optional parameter *k* specifies the number of nearest neighbors to use when calculating the graph neighborhood radius *r*_0_. The value of *k*, if set, is the minimum (respectively maximum) number of neighbors that most clustered (respectively unclustered) localizations should have. It should be smaller than the number of localizations in a typical cluster, but preferably larger than the estimated number of times that a typical single molecule might blink. These values can be estimated by visual inspection of localization clusters within cell boundaries and on the coverslip outside of cells. Increasing *k*, and consequently *r*_0_, can influence the exact placement of cluster boundaries, and hence cluster quantification, by allowing more low-density localizations on the periphery of clusters to be included in the clusters. This highlights the inherent ambiguity in clustering problems, which results from the lack of a clear definition of a cluster. We recommend values of *k* between 10 and 20 for most data. Alternatively, StormGraph can determine *r*_0_ heuristically without any user input (i.e. without *k*), but this approach is designed for data with very few localizations dispersed in between clusters. Set *k* = 0 in the software to use this mode of StormGraph. We advise that this mode should only be used if it is clear *a priori* that at least two thirds of localizations in each ROI belong to clusters.

The parameter *α* controls the node-degree threshold used to identify and remove unclustered nodes prior to clustering. For data that does not suffer from overcounting of molecules, or for which overcounting has already been corrected, *α* is effectively the maximum false positive rate (FPR) for classifying localizations as clustered if all localizations in a random distribution should be classified as unclustered. It can be regarded as a per-localization significance level. When overcounting is present in the data, the FPR may be greater than *α*. Nevertheless, for any given *α* < 1, StormGraph takes steps to minimize the FPR as far as possible. Hence, we suggest setting *α* as the maximum fraction of localizations that the user would accept as being clustered if they were completely randomly distributed. For most applications, we recommend *α* = 0.05, the default value. Larger values of *α* might be suitable if the user is already confident that the localizations are strongly clustered but there is large variation in the density of clusters. For example, *α* = 0.5 would simply demand that clusters are at least as dense as the average density of a random distribution, but this could result in as many as 50% of localizations in a random distribution qualifying as clustered. Alternatively, the user can choose to skip the thresholding step and instead allow all localizations to be possibly assigned to clusters by setting *α* = 1, which ultimately removes all use of *α* and *k* from the StormGraph algorithm.

Finally, the user can optionally set the minimum number of localizations that a cluster must contain, MinCluSize. One possible strategy for setting its value is to investigate clusters of localizations in background regions outside of cells, which are likely to be due to individual fluorescent labels stuck to the coverslip, and assess how many localizations are typical of these apparent clusters. However, because StormGraph provides an option to use localization uncertainties to identify and reclassify localization clusters that could have arisen just from overcounting of single molecules, clusters that could be due to single molecules can be automatically removed from analysis without the need for a minimum cluster size parameter. Note that StormGraph requires all clusters to contain at least three localizations, even if MinCluSize is not set.

### Computational approximations in StormGraph

In order to improve computational efficiency, StormGraph includes some computational approximations. Firstly, neighborhood searches about each node are performed using the MATLAB function “rangesearch”, which uses a *k*-d tree, as this is faster than computing distances between all pairs of nodes. Without uncertainties in localization positions, rangesearch is implemented with a search radius of *r*_0_. However, when Monte Carlo simulations are used to perturb localization positions using their uncertainties, it is inefficient to perform rangesearch for every simulation. Instead, StormGraph performs rangesearch just once, using an expanded search radius, to identify candidate edges for the graph. It then calculates expected edge weights only for the candidate edges. Since the computational time for rangesearch increases as the search radius increases, we chose (*r*_0_ + 6 × mean localization uncertainty) as the expanded search radius because most pairs of nodes separated by distances greater than this would have only negligible or zero edge weights anyway. Increasing the search radius further would not only make rangesearch slower, but it could also add more edges to the graph and consequently increase the computational cost of community detection, even though the additional edges would be mostly negligible.

Secondly, StormGraph limits nodes to having no more than 500 neighbors in the graph. This is to prevent extremely dense, large clusters from dramatically slowing down community detection, since the computational time required by Infomap scales with the number of edges in the graph. In practice, for reasonably chosen values of *k*, e.g. in the range from 10 to 20, and for *r*_0_ values determined heuristically, very few nodes, if any, in most datasets should have this many neighbors.

Lastly, we note that StormGraph is not deterministic, meaning that it can give slightly different results each time that it is run. This is for two reasons. The first reason is because StormGraph uses Infomap or the Louvain method to perform community detection. Infomap seeks to optimize the map equation and the Louvain method seeks to optimize modularity. In both cases, the full optimization problem is NP-hard. Therefore, both methods take a greedy approach to the optimization, which generally finds a local, but not necessarily global, optimum. They then select the best optimum from multiple iterations started from random initiations. In StormGraph, the default number of iterations used for finding the final cluster hierarchy is 50. Results can be improved at the expense of increasing computational time by increasing the number of iterations. Conversely, computational time can be reduced at the expense of cluster accuracy by decreasing the number of iterations. The second reason for slight variability in results is the use of Monte Carlo simulations by StormGraph. This variability can be decreased, again at the expense of increasing computational cost, by increasing the number of Monte Carlo simulations.

The non-deterministic nature of StormGraph is only a minor drawback, as variability in clustering results for a single dataset is small. To demonstrate this, we repeatedly applied StormGraph using identical settings to a heterogeneous dSTORM ROI containing visually ambiguous clusters. We did this in both 2D and 3D and for both the automatic and kNN methods for determining *r*_0_, each time generating 11 StormGraph repeats. We then assessed the similarity of cluster assignments from each of the last 10 repeats to the first one using NMI, which can range from 0 to 1. We always achieved NMI > 0.94, indicating very high similarity (Figure S3).

### Simulating SMLM data in 2D and 3D

In both 2D and 3D, except for the simulations used to test the Bayesian clustering method (29), we distributed 3,000 molecules into circular nanoclusters with a fixed radius, *r*. These molecules were assigned to nanoclusters uniformly at random with a fixed average molecular density, *ρ*. Each molecule was assigned uncertainties, which were sampled randomly from a real dSTORM dataset, in its *x*-, *y*- and (for 3D) *z*-coordinates and a number of blinks, which was drawn from a geometric distribution (55) supported on {1, 2, 3,…} with success probability parameter λ. Within each nanocluster, molecules were distributed uniformly at random, and for each molecule the observed localizations (blinks) were drawn from a normal distribution with mean equal to the molecule’s position and standard deviations equal to the uncertainties assigned to the molecule. Every observed localization was assigned the same uncertainties as its associated molecule. The total number of nanoclusters, *N*_nano_, was determined by the total number of molecules in clusters (3,000) and the average density, *ρ*, of molecules within clusters.

The nanoclusters were positioned inside a 2 μm × 2 μm ROI in 2D or a 2 μm × 2 μm × 1 μm ROI in 3D such that some existed as isolated nanoclusters and others were randomly aggregated into larger clusters according to the following process, which was adapted from a Dirichlet process: for *i* from 1 to *N*_nano_, draw a random number from the uniform distribution on [0,1]; if it is less than or equal to ((*p* + 10)/(*p* + *i* – 1))^*q*^ for positive integers *p* and *q*, then place the *i*^th^ nanocluster away from existing clusters; otherwise, add the *i*^th^ nanocluster to a randomly selected existing cluster, excluding the first 10 nanoclusters that were placed. If a nanocluster was added to an existing cluster, it was placed such that its centre was exactly a distance 2*r* from the centre of another nanocluster in the same aggregate cluster, and without overlapping with any other existing nanoclusters in the aggregate cluster.

This process ensures that there are at least 10 isolated nanoclusters and a variable number of larger aggregate clusters of variable size, thus creating heterogeneous clusters. The heterogeneity is controlled by the parameters *p* and *q*. In our simulations, we fixed *p* = 5 and varied *q* from 1 to 5, with larger values of q resulting in larger (and fewer) cluster aggregates. Outside of the clusters, we added molecules uniformly at random at a specified average density, and the number and positions of observed localizations corresponding to each of these background molecules were drawn from geometric and normal distributions respectively, as described for the in-cluster localizations.

If the simulations were performed in 3D, points were then randomly removed such that the probability of a localization being observed in the final simulated data decayed according to a Gaussian profile as the axial distance from a central focal plane increased. This was to imitate the realistic scenario for most 3D SMLM techniques in which fluorescent blink events are more likely to be collected and localized the closer they are to the focal plane.

We generated 64 2D datasets with multiple blinking of molecules (e.g. Figures 2a–c and S7) by varying the following parameters: (1) the radius of the nanoclusters (20 nm, 30 nm or 50 nm), (2) the density of clustered molecules (0.01 nm^−2^ or 0.02 nm^−2^), (3) the density of the random molecules (1%, 5%, 10%, 20% or 40% of the within-cluster molecular density), (4) the average number of blinks per molecule (4/3, 2 or 4; these values provide examples ranging from cases in which most molecules blink only once to cases where the molecules could be bivalent and labeled by fluorophores that blink on average twice, which is typical for the photoactivatable fluorophore mEos2 (14; 56)), and (5) the propensity for nanoclusters to coalesce into larger aggregate clusters (parameter *q*).

We generated 130 3D datasets analogously but using within-cluster molecular densities of 1 × 10^−4^ nm^−3^ and 2 × 10^−4^ nm^−3^. In 3D, we used nanoclusters of radii 30 nm and 50 nm, and we used densities of random, unclustered molecules equal to 1%, 5%, 10% or 20% of the within-cluster molecular density. At 20%, clusters were barely visible in 2D projections of the simulated 3D data onto the *xy*-plane.

Simulations of 1 μm × 1 μm ROIs for comparison of StormGraph to the Bayesian method of Rubin-Delanchy et al. (29) followed the above methods for simulating 2D ROIs but with the following modifications. The total number of molecules belonging to clusters was reduced from 3,000 to 1,000. The minimum number of isolated nanoclusters guaranteed to not form larger aggregates was reduced from 10 to 3. Circular nanoclusters had radius *r* = 30 nm and average molecular density *ρ* = 0.01 nm^−2^ in all simulations. The values used for the average number of blinks per molecule were 4/3 and 2, and the values used for the parameter *q* were 1, 3, and 5. We simulated a total of 30 ROIs that were 1 μm × 1 μm.

### Running ClusterViSu on simulated data

The ClusterViSu algorithm consists of running a series of two functions provided as part of its source code, specifically the functions “VoronoiMonteCarlo” and “VoronoiSegmentation”. However, the authors did not provide a script for running ClusterViSu. Hence, for users with zero programming expertise, it can only be run using a graphical user interface that requires each file to be loaded and analyzed separately. Also, ClusterViSu outputs the bounding polygon for each detected cluster but not the actual cluster assignments of the localizations, which we needed to compute NMI and mean F-measure scores for assessing the performance of cluster assignment. Therefore, we wrote our own custom MATLAB script (available upon request) to run and batch process ClusterViSu from its source code and subsequently determine the cluster assignments of the localizations. In addition, ClusterViSu prefers input ROIs to be at least 18 μm × 18 μm, so we rescaled our 2 μm × 2 μm simulated data by a factor of 9, which drastically improved ClusterViSu’s performance, at least in terms of computational time.

Furthermore, we only included ClusterViSu results for simulated datasets on which ClusterViSu analysis completed in under 2 hours. This resulted in 15 out of 64 simulated datasets being excluded from our summary of test results for ClusterViSu, but these 15 datasets were still included for assessing StormGraph and DBSCAN. However, these 15 datasets were excluded in Figures 2d and S4a(iii), where NMI or mean F-measure results for StormGraph and DBSCAN are shown as a ratio to the NMI or mean F-measure results for ClusterViSu.

### Functionalization of glass coverslips for cell adherence

Glass coverslips were cleaned and functionalized as previously described (57). Briefly, acid-cleaned glass coverslips (Marienfeld #1.5H, 18 mm × 18 mm; catalogue #0107032, Lauda-Königshofen, Germany) were incubated with 0.01% poly-L-lysine (Sigma-Aldrich; catalogue #P4707) or 0.25 μg/cm^3^ of the non-stimulatory M5/114 anti-MHCII monoclonal antibody (Millipore; catalogue #MABF33) or 2 μg/cm^2^ fibronectin (Sigma Aldrich; catalogue #F4759) for at least 3 h at 37 °C. The slides were then washed with phosphate-buffered saline (PBS) prior to being used for experiments.

### Monovalent Fab fragments and antibodies

The anti-mouse-Igκ antibody for clustering BCRs was purchased from Southern Biotech (Birmingham, AL; catalogue #1050-01). AF647-conjugated anti-mouse-IgM Fab fragments (catalogue #115-607-020) and AF647-conjugated anti-human-IgM Fab fragments (catalogue #109-607-043) were from Jackson ImmunoResearch Laboratories (West Grove, PA). All Fab fragments were routinely tested for aggregation using dynamic light scattering (Zetasizer Nano) and unimodal size distributions were observed. Anti-LAMP-1 antibody was purchased from Abcam (catalogue #ab24170). AF647-conjugated goat anti-mouse-IgG (catalogue #A21236) and AF647-conjugated goat anti-rabbit-IgG (catalogue #A21244) were purchased from ThermoFisher Scientific. Goat anti-mouse-IgG (Jackson ImmunoResearch Laboratories; catalogue #115-005-008) and goat anti-rabbit-IgG (Jackson ImmunoResearch Laboratories; catalogue #111-001-008) were conjugated to Cy3B using a Pierce antibody conjugation kit (catalogue #44985).

### Cell labeling for dSTORM

#### (1) Murine splenic B cells

Animal protocols were approved by the University of British Columbia and all animal experiments were carried out in accordance with institutional regulations. Splenic B cells were obtained from 6- to 10-week old C57BL/6 mice (Jackson Laboratory) of either sex using a B-cell isolation kit (Stemcell Technologies; catalogue #19854) to deplete non-B cells. To induce IgM-BCR clustering, 5 × 10^6^ *ex vivo* splenic B cells/mL were stimulated with 20 μg/mL anti-Igκ in PBS for 10 min at 37 °C. A similar volume of PBS was added to control samples (resting B cells). All subsequent procedures were performed at 4 °C. Cells were washed three times with ice-cold PBS, and IgM-BCRs on the cell surface were labeled using AF647-conjugated monovalent anti-mouse-IgM Fab fragments for 15 min. These Fab fragments bind to the constant region of the μ heavy chain of IgM-BCRs, which is distinct from sites on the IgM-BCR that the anti-Igκ treatment antibody binds to. Following multiple PBS washes, cells were settled onto pre-cooled anti-MHCII-functionalized coverslips for 10 min and subsequently fixed with PBS containing 4% paraformaldehyde and 0.2% glutaraldehyde for 90 min. The coverslips were washed thoroughly with PBS and fiducial markers (100 nm diameter; ThermoFisher Scientific, catalogue #F8799) were allowed to settle onto the coverslip overnight at 4 °C. Unbound fiducial markers were removed by PBS washes and the stuck particles were used for real-time drift stabilization (58).

#### (2) Human and murine B-lymphoma cell lines

A20 and BJAB B-lymphoma cells were obtained from American Type Culture Collection (ATCC). HBL-1 cells were obtained from Dr. Izidore S. Lossos, Sylvester Comprehensive Cancer Center, University of Miami (Miami, FL). TMD8 cells were a gift from Dr. Neetu Gupta, Lerner Research Institute, Cleveland Clinic (Cleveland, OH). All B-cell lines were cultured in RPMI-1640 (Life Technologies; catalogue #21870-076), supplemented with 10% heat-inactivated fetal bovine serum, 2 mM L-glutamine, 50 μM β-mercaptoethanol, 1 mM sodium pyruvate, 50 U/mL penicillin, and 50 μg/mL streptomycin (complete medium). All cell lines were authenticated by STR DNA profile analysis.

All staining procedures were performed at 4 °C. Cell-surface IgM-BCRs on BJAB, HBL-1 and TMD8 cells were labeled using AF647-conjugated anti-human-IgM Fab fragments for 15 min. Cell-surface IgG-BCRs on A20 cells (ATCC) were labeled using both AF647-conjugated anti-mouse-IgG and Cy3B-conjugated anti-mouse-IgG at 1:1 stoichiometry for 15 min. Fc receptors on A20 cells were blocked prior to staining using the 2.4G2 rat anti-Fcγ receptor monoclonal antibody. Cells were washed in PBS and subsequently fixed with ice-cold PBS containing 4% paraformaldehyde and 0.2% glutaraldehyde for 60 min. Following multiple PBS washes, the cells were settled onto pre-cooled poly-L-lysine-coated coverslips for 15 min and subsequently fixed again for 30 min. The coverslips were washed thoroughly with PBS and fiducial markers were added and incubated overnight at 4 °C.

#### (3) B16 melanoma cell lines

B16F1 melanoma cells (ATCC) were grown in RPMI-1640 complete medium. Approximately 3 × 10^4^ cells were seeded on fibronectin-coated coverslips for 1 h and fixed with PBS containing 4% paraformaldehyde for 30 min. Cells were permeabilized with 0.1% Triton X-100 for 10 min, washed with PBS, and incubated for 30 min at room temperature (RT) with Image-IT FX Signal Enhancer (Life Technologies, catalogue #I36933) to neutralize surface charge. Cells were washed briefly in PBS and then incubated with BlockAid blocking solution (Life Technologies; catalogue #B10710) for 1 h at RT. The cells were incubated with anti-LAMP-1 antibody (diluted in BlockAid) for 4 h at RT. Following PBS washes, cells were incubated with both AF647-conjugated anti-rabbit-IgG and Cy3B-conjugated anti-rabbit-IgG at 1:1 stoichiometry for 90 min. Cells were washed in PBS and subsequently fixed again with 4% paraformaldehyde for 10 min. The coverslips were washed thoroughly with PBS and fiducial markers were added and incubated overnight at 4 °C.

### dSTORM

Imaging was performed using a custom-built microscope with a sample drift-stabilization system that has been described previously (58; 59). Briefly, three lasers were used in the excitation path. These were a 639 nm laser (Genesis MX639, Coherent) for exciting the AF647, a 532 nm laser (Laser quantum, Opus) for exciting the photo-switchable Cy3B, and a 405 nm laser (LRD 0405, Laserglow Technologies) for reactivating the AF647 and Cy3B. All three lasers were coupled into an inverted microscope equipped with an apochromatic TIRF oil-immersion objective lens (60x; NA 1.49; Nikon). The emission fluorescence was separated using appropriate dichroic mirrors and filters (Semrock) (58; 59), and detected by EM-CCD cameras (Ixon, Andor). A feedback loop was employed to lock the position of the sample during image acquisition using immobile fiducial markers. Sample drift was controlled to be less than 1 nm laterally and 2.5 nm axially.

### dSTORM image acquisition and reconstruction

Imaging was performed in an oxygen-scavenging GLOX-thiol buffer consisting of 50 mM Tris-HCl, pH 8.0, 10 mM NaCl, 0.5 mg/ml glucose oxidase, 40 μg/ml catalase, 10% (w/v) glucose and 140 mM 2-mercaptoethanol (60). The coverslip with attached cells was mounted onto a depression slide filled with imaging buffer and sealed with Twinsil two-component silicone-glue (Picodent; catalogue #13001000).

For SMLM imaging, a laser power density of 1 kW/cm^2^ for the 639 nm and 532 nm lasers was used to activate the AF647 and Cy3B, respectively. For each sample, 4 × 10^4^ images were acquired for each color channel at 50 Hz. Localization coordinates and their associated uncertainties were computationally determined simultaneously by fitting a function to the intensity profile of each fluorescence event using MATLAB (Figure S13), as described previously (59). Expressed as standard deviations, lateral uncertainties were typically < 10 nm while axial uncertainties were typically < 40 nm (Figure S13).

For two-color SMLM, image acquisition was performed sequentially for each color with AF647 imaged first to prevent photobleaching by the Cy3B excitation laser. Two-color SMLM images were acquired using a beam splitter with appropriate filters to direct each signal to one of two independent cameras. Alignment of these two colors was carried out using ~4 × 10^4^ images of fluorescent beads simultaneously recorded at various positions to find an optimal geometric transformation. The resulting color-alignment error is ~10 nm root mean squared.

## Acknowledgements

We thank Alejandra Herrera-Reyes for helpful discussions and code, Ki Woong Sung for preliminary computational work, Dr. Vivian Qian Liu for assistance with dSTORM data fitting, Dr. Neetu Gupta for TMD8 cells, Dr. Izidore S. Lossos for HBL-1 cells, and Dr. David R.L. Scriven for helpful discussion. This work was supported by funding from the Natural Science and Engineering Research Council of Canada (Discovery Grant RGPIN-2015-04611 to DC, an Undergraduate Student Research Award to DWZ, Discovery Grant RGPIN-2014-03581 to KCC, and Discovery Grant RGPIN-2017-04862 to MRG), the Canadian Cancer Society Research Institute (Innovation Grant 704254 to DC and MRG), the Canadian Institutes of Health Research (PJT-19426 to MRG), and Canada Foundation for Innovation (to KCC).

## Author contributions

JMS conceived the project, developed and tested the StormGraph algorithm and software, proposed experiments, performed data analysis, produced figures, and wrote the manuscript. LA proposed and performed experiments, wrote experimental methods, produced figures, and provided essential feedback about the algorithm and software. DWZ assisted with software development, simulation of data, and algorithm testing. RT built the dSTORM microscope and assisted with dSTORM data fitting and processing. KCC provided code for fitting dSTORM localizations and aligning two-color dSTORM data. MRG and DC supervised the project and wrote the manuscript. All authors approved the final manuscript.

