## Supplementary information file for "StormGraph: A graph-based algorithm for quantitative clustering analysis of diverse single-molecule localization microscopy data"

#### Supplementary Notes

##### Supplementary Note 1: Resolution limits of community detection algorithms

The Louvain method is subject to a resolution limit (1), whereby the minimum size of detected clusters increases as the size of the graph increases. However, each implementation of the Louvain method generates a hierarchical clustering, with the resolution limit applying to the highest (most coarse) level. To minimize the effect of the resolution limit on the preliminary clustering performed by StormGraph during node degree thresholding when localization uncertainties are unknown, the lowest (finest) level of the hierarchy is used (matching the approach taken by Emmons et al. in their comparison of community detection algorithms (2)). The two-level Infomap algorithm is also subject to a resolution limit, but it is orders of magnitude smaller than that of the Louvain method (3) and therefore not a major concern. For best results, the lowest level of a hierarchical clustering optimized by the multi-level Infomap method, which is not subject to a resolution limit (3), could be used instead, but for computational efficiency we use the faster two-level method during the preliminary clustering step.

### Supplementary Tables

#### Table S1

| *Algorithm:* | SR-Tesseler | Cluster-ViSu | DBSCAN | OPTICS | FOCAL | Bayesian method | Storm-Graph |
|---|---|---|---|---|---|---|---|
| **References** | (4) | (5; 6) | (7) | (8) | (9) | (10; 11) | N/A |
| **Mathematical approach** | Voronoi tessellations | Voronoi tessellations | Density-based | Density-based | Pixel-based | Bayesian, model-based | Graph-based |
| **Robust to ROI-to-ROI heterogeneity** | yes | yes | no | no* | yes | no | yes |
| **3D capability** | # | yes | yes | yes | no | yes | yes |
| **Adapts to axial variation in 3D SMLM data** | N/A | yes | no | yes | N/A | yes** | yes |
| **Suitable for arbitrary cluster shapes and sizes** | yes | yes | yes | yes | yes*** | no | yes |
| **Accounts for localization errors** | no | no | no | no | ## | yes | yes |
| **Includes colocalization analysis** | # | yes | no | no | no | no | yes |
| **Hierarchical clustering** | no | no | no | yes | no | no | yes |
| **Automatically generates single-level clustering** | yes | yes | yes | no | yes | yes | yes |
| **Results are deterministic** | yes | no**** | yes | yes | yes | yes | no |
| **Number of inputs controlled by user** | $\geq 1$ | 0–1**** | 2 | 2 | 0–1 | 5 | 1–3 |

**Table S1. Comparison of features of StormGraph and other clustering algorithms.**
### During revision of this manuscript, authors of SR-Tesseler published a related method called Coloc-Tesseler (12) that is designed specifically for colocalization analysis of 2D and 3D SMLM data.
#### Uses a single global estimate of localization uncertainty to define a pixel size, but does not account for each individual localization's specific positional uncertainty.
* Requires user-specification of the same two density-based parameters as DBSCAN, but theoretically the search radius $\varepsilon$ can be made arbitrarily large, unlike for DBSCAN. This means that, theoretically, OPTICS can operate independently of the global localization density (i.e. it can be implemented with parameters that do not depend on the density). However, this would result in all localizations being grouped together in a single cluster at the most coarse level of the cluster hierarchy and would be computationally inefficient. Therefore, in practice, the parameter $\varepsilon$ should be set.
** Technically possible by modifying the mathematical model used by the algorithm, but this is non-trivial.
*** Yes, except that clusters will be composed of square pixels.
**** Assumes that ClusterViSu is operated, as intended, using Monte Carlo simulations to automatically determine its threshold Voronoi cell area (or volume) needed to define clusters.

### Supplementary Figures

#### Figure S1

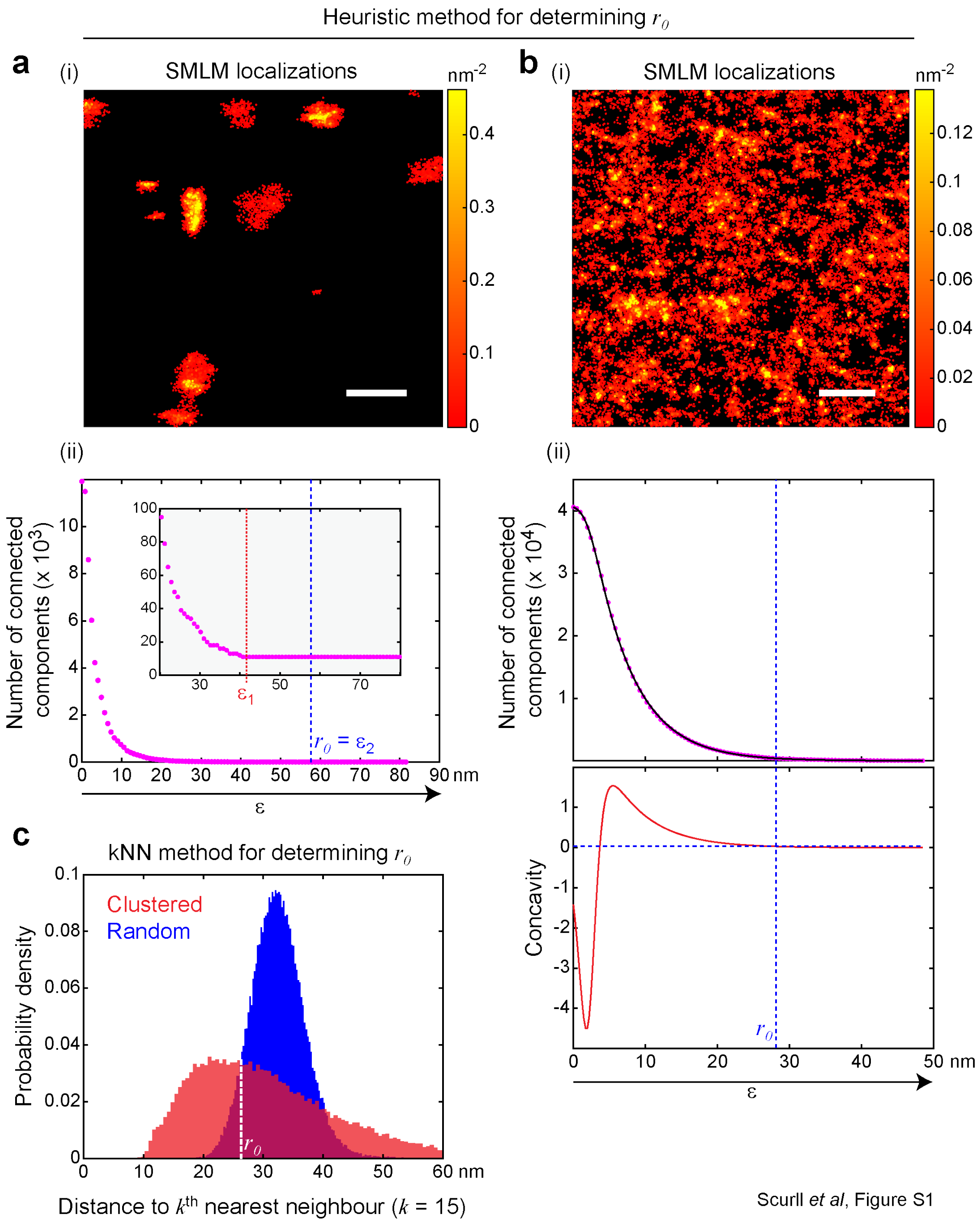


**Figure S1. The two available methods for setting the search radius $r_0$ in StormGraph. (a)** The heuristic elbow method for setting $r_0$ when there is an innate number of clusters in the data. (i) Example

SMLM localizations (adapted from real dSTORM data of IgG-BCRs on an A20 murine B cell) with well defined clusters and minimal dispersed localizations between clusters. Color bar = density ($nm^{-2}$), scale bar = 500 nm. (ii) Number of connected components in the $\varepsilon$-neighborhood graph vs $\varepsilon$ for the data in (i). StormGraph sets $r_0 = \varepsilon_2$, where an $\varepsilon_2$-neighborhood has twice the area/volume of an $\varepsilon_1$-neighborhood, if the number of connected components remains constant for $\varepsilon_1 \leq \varepsilon \leq \varepsilon_2$. Inset: enlarged region of plot. **(b)** (i) Example SMLM localizations with poorly defined clusters and many dispersed localizations between clusters. Color bar = density ($nm^{-2}$), scale bar = 500 nm. (ii) The heuristic elbow method for setting $r_0$ for the data in (i). A function $f(\varepsilon)$ is fit to number of connected components vs $\varepsilon$ for the $\varepsilon$-neighborhood graph (top). The concavity $f''(\varepsilon)$ of this curve is then computed (bottom), and the value of $r_0$ (vertical dashed line) is set as the value of $\varepsilon$ at which $f''(\varepsilon)$ returns to 2% (horizontal dashed line) of its maximum value. **(c)** The k-nearest neighbor (kNN) method for setting $r_0$ (illustrated for the data in panel (b)). A histogram of distances from nodes to their $k^{\text{th}}$ nearest neighbors is obtained for both the clustered SMLM data (red) and a random distribution of points with the same average density (blue), where $k$ is a user-specified parameter. For data with known localization uncertainties, the 95% confidence level from Monte Carlo simulations is used for the $k^{\text{th}}$ nearest neighbor distance for each localization. The value of $r_0$ (vertical white dashed line) is set as the distance at which the two histograms first intersect.

### Figure S2

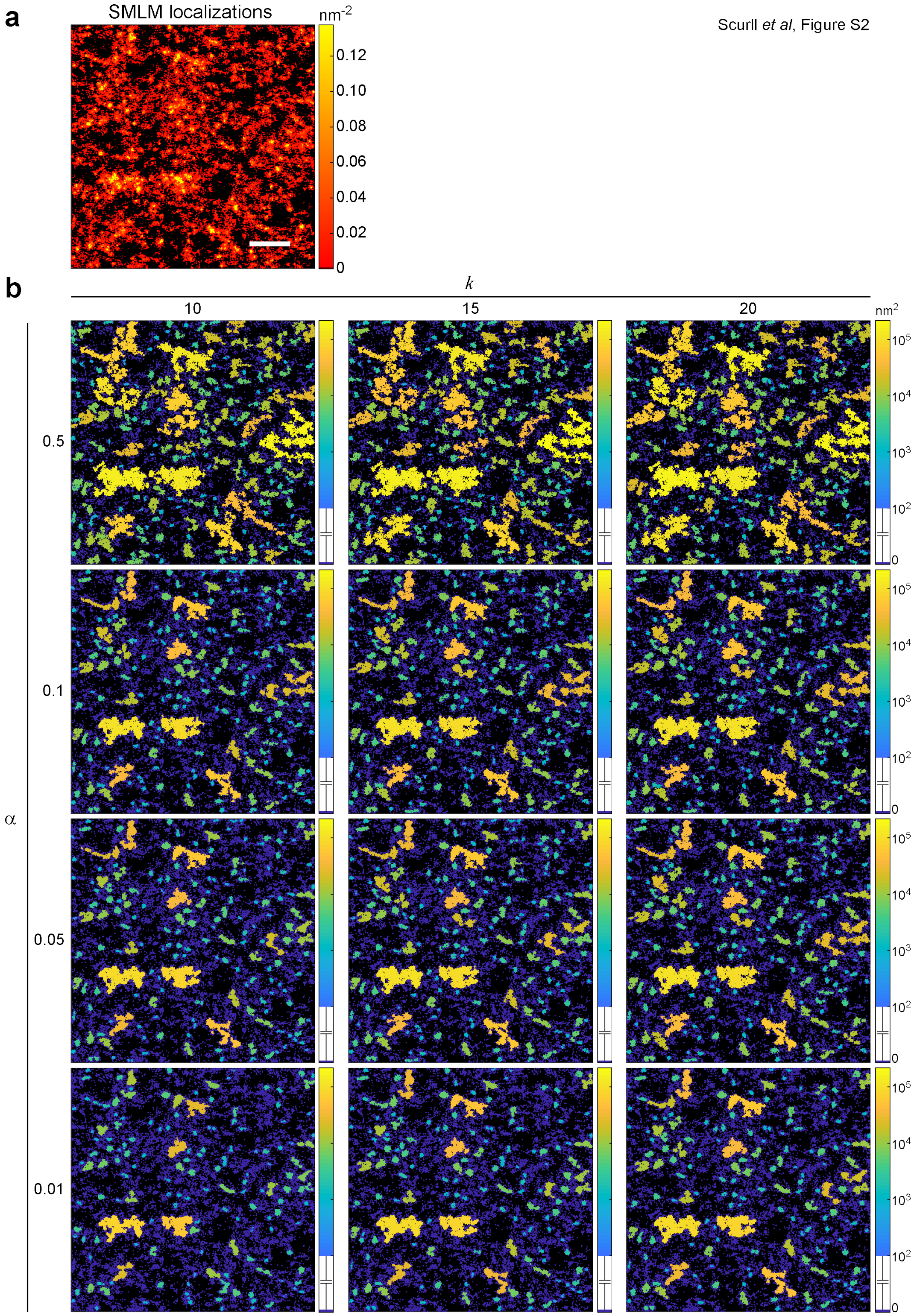

a
SMLM localizations
nm-2
0.12
0.10
0.08
0.06
0.04
0.02
0
Scurll et al, Figure S2
b
k
10
15
20
nm2
α
0.5
0.1
0.05
0.01
10^5
10^4
10^3
10^2
0

**Figure S2. Effect of varying StormGraph's parameters $k$ and $\alpha$ for an example SMLM ROI.** **(a)** An ROI containing localizations of Alexa Fluor 647-labeled cell-surface B-cell antigen receptors of IgM isotype on a TMD8 diffuse large B-cell lymphoma cell imaged by dSTORM. Scale bar = 500 nm, color bar = density ($nm^{-2}$). **(b)** Clusters identified by StormGraph using different values of the parameters $k$ and $\alpha$. In all cases, StormGraph was implemented in 2D, using localization uncertainties, and with a minimum cluster size of 5 localizations. Clusters are colored according to their areas ($nm^2$) calculated by StormGraph. A cluster area of 0 (dark blue) indicates that a localization was classified as unclustered (i.e. not assigned to any cluster).

#### Figure S3

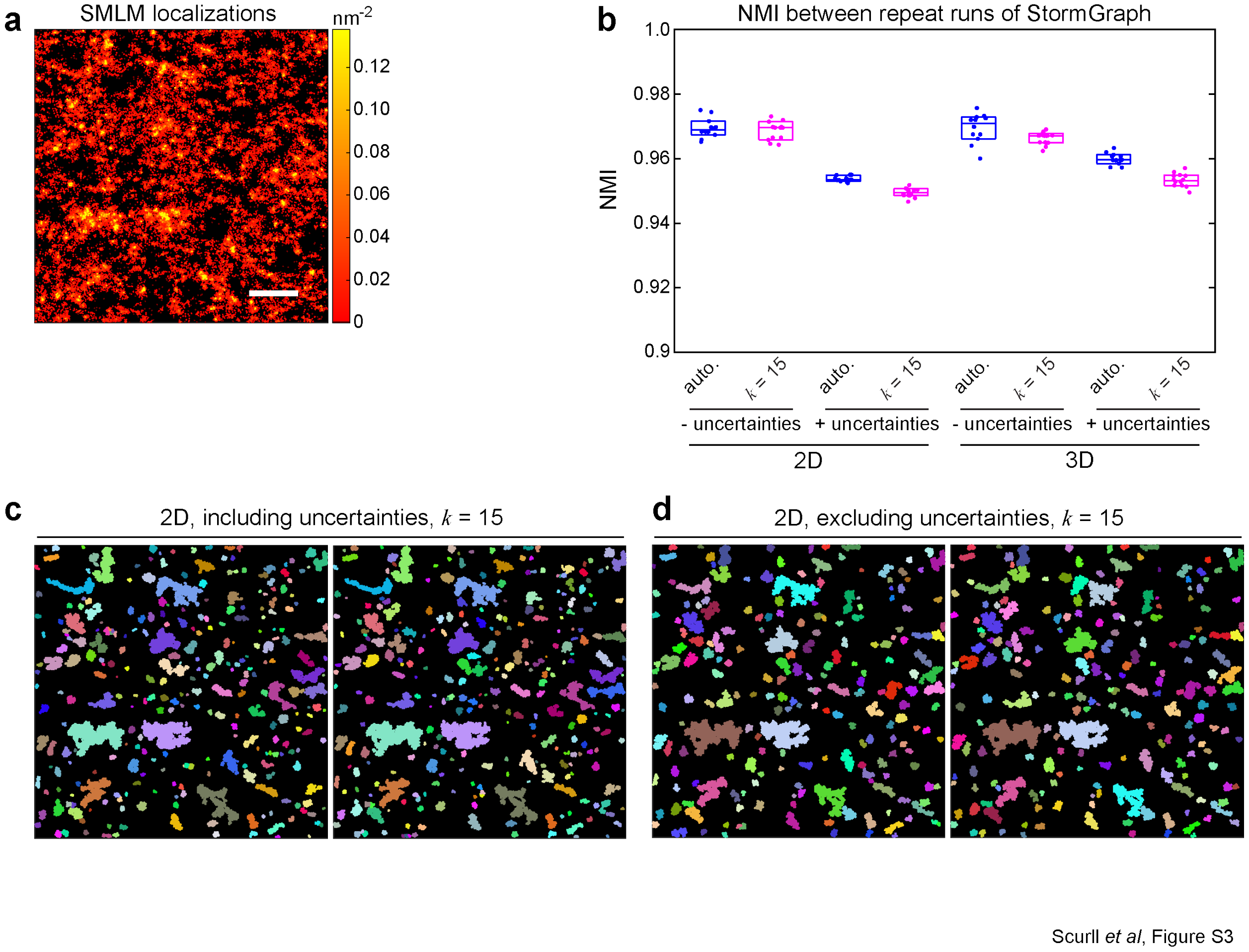


**Figure S3. Identical repeat runs of StormGraph yield highly similar results. (a)** 2D projection of a 3D ROI containing heterogeneous localizations of AF647-labeled cell-surface IgM-BCRs on a TMD8 B cell imaged by dSTORM. Scale bar = 500 nm, color bar = density ($nm^{-2}$). **(b)** Normalized mutual information (NMI) measuring the similarity between single-level cluster assignments of localizations by multiple identical runs of StormGraph. For each of the indicated StormGraph settings, StormGraph was run identically 11 times on the data in (a). The NMI values in the figure score the similarity of the cluster assignments of localizations by the first run of StormGraph to the cluster assignments of localizations by each of the 10 subsequent identical runs of StormGraph. StormGraph was implemented in 2D or 3D either not using (-) or using (+) localization uncertainties. The value of $r_0$ was determined using either the heuristic method (auto.) or the kNN method with $k = 15$. NMI values can range from 0 to 1, with NMI = 1 corresponding to perfect similarity. Boxes show medians and interguartile ranges. **(c–d)** Single-level clusters (distinguished by colors) identified by one run of StormGraph (left) and the *least similar* results of 10 subsequent identical runs of StormGraph using the indicated settings.

### Figure S4

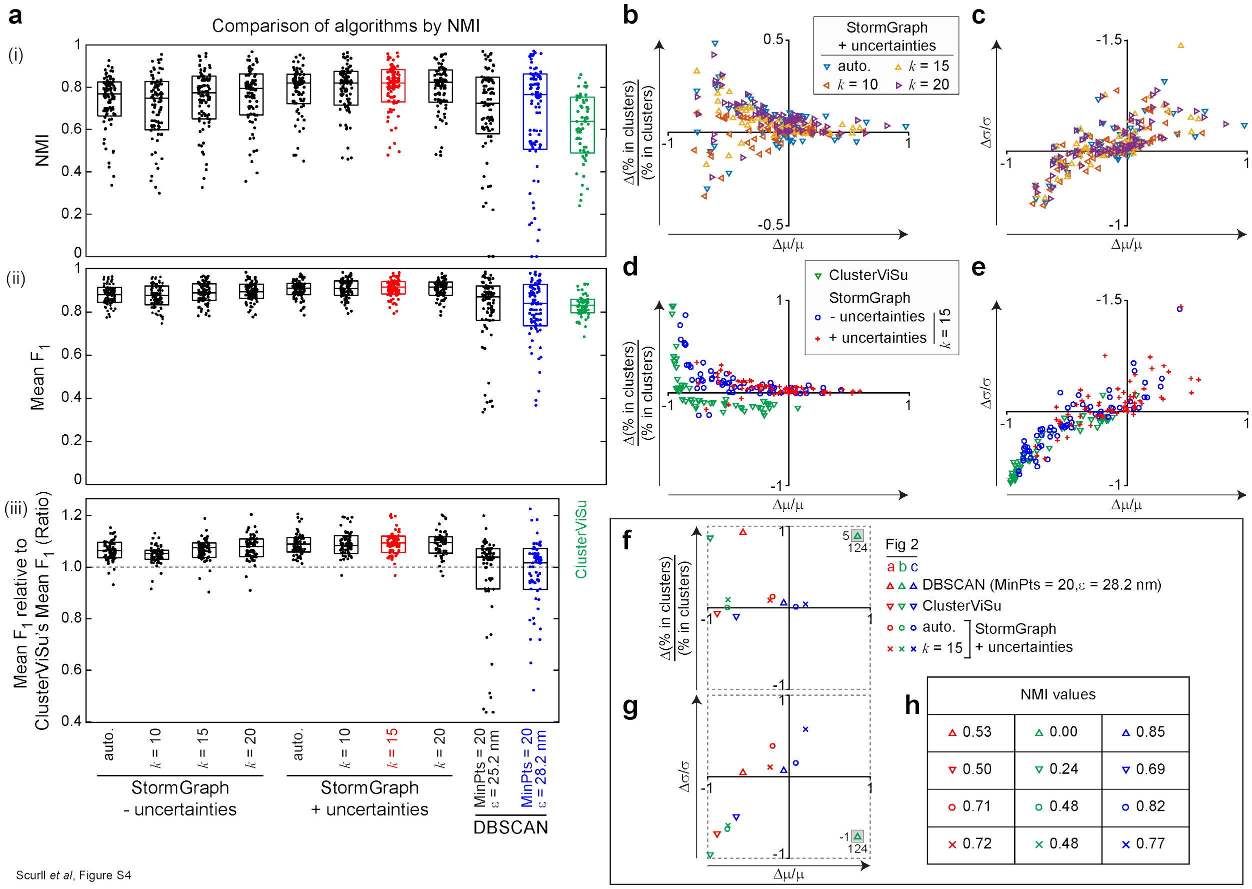


**Figure S4. StormGraph is robust and more accurate than ClusterViSu and DBSCAN when analyzing simulated data with known ground-truth clusters. (a)** Accuracy of assigning data points to clusters as assessed by (i) normalized mutual information (NMI) and (ii) mean F-measure ($F_1$); 1 = 100% agreement with ground-truth. A total of 64 simulated datasets were analyzed using StormGraph, ClusterViSu, and DBSCAN. StormGraph was run either with (+) or without (-) localization uncertainties, always with $\alpha = 0.05$ and MinCluSize = 5. The value of $r_0$ used by StormGraph was determined using the heuristic method (auto.) or the kNN method with $k = 10$, 15 or 20. DBSCAN was implemented using 16 different selections of its two parameters, MinPts and $\varepsilon$, of which the two best-performing are shown here. ClusterViSu results are only shown for the 49 datasets on which the analysis was completed in under 2 h. Boxes show medians and interquartile ranges. The mean F-measure results in panel (b) were additionally normalized to ClusterViSu's performance for each of the 49 simulated datasets for which analysis by ClusterViSu was completed in under 2 h (iii). StormGraph was consistently more accurate than ClusterViSu at assigning points to clusters, as indicated by ratios $> 1$. The colors used in (a) match the colors used in Figure 2(d–f) for reference only. **(b–g)** Cluster quantification errors relative to ground truth. The fractional error in the percentage of localizations assigned to clusters (b, d, f) and the fractional error in the standard deviation, $\sigma$, of number of localizations per cluster (c, e, g) are plotted *versus* the fractional error in the mean number of localizations per cluster, $\mu$, for each of the 64 simulated ROIs for StormGraph and DBSCAN. For ClusterViSu, the errors are plotted for the 49 ROIs for which analysis was completed in under 2 h. N.b. $\pm 1$ means $\pm 100\%$ error. Panels (b–c) compare quantification errors by StormGraph using different values of $k$ or the heuristic (auto.) method for determination of $r_0$. Panels (d–e) compare

ClusterViSu and StormGraph with and without using localization uncertainties for $k = 15$. Panels (f–g) show quantification errors by StormGraph, ClusterViSu, and DBSCAN for the specific example simulated ROIs shown in Figure 2(a–c). The shaded boxes show axes values for scatter points that lie outside the displayed axes ranges. **(h)** Normalized mutual information (NMI) values for the specific example simulated ROIs shown in Figure 2(a–c).

#### Figure S5

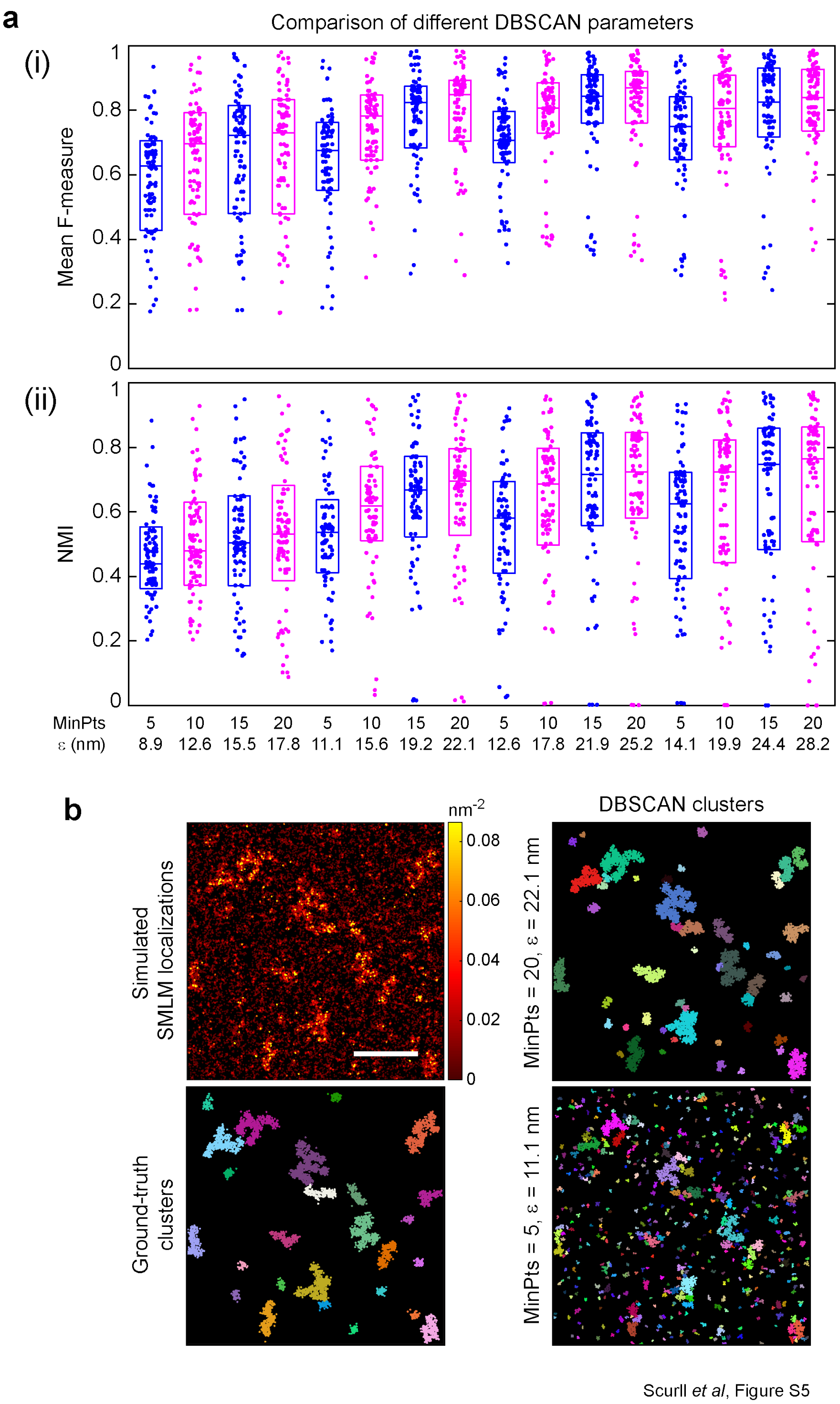


**Figure S5. DBSCAN is sensitive to the choice of parameters, and no single choice of parameters is suitable for batch processing cluster analysis when localization density is variable**

**between datasets. (a)** Accuracy of cluster assignments by DBSCAN, using different values of the parameters MinPts and $\varepsilon$, of localizations in the 64 simulated datasets used for Figure 2b. Accuracy was measured by (i) mean F-measure and (ii) normalized mutual information (NMI). A total of 16 parameter pairs corresponding to four different threshold densities (0.020, 0.013, 0.010 and 0.008 $nm^{-2}$) were tested. Boxes show medians and interquartile ranges. **(b)** A visual demonstration that two different DBSCAN parameter choices corresponding to the same threshold density produce very different clustering results. Top left: the simulated dataset example used in Figure 2a; color bar = density ($nm^{-2}$); scale bar = 500 nm. Bottom left: ground-truth clusters present in the example simulated dataset; colors distinguish distinct clusters. Right panels: clusters identified by DBSCAN using two different parameter pairs, both of which correspond to a threshold density of 0.013 $nm^{-2}$.

#### Figure S6

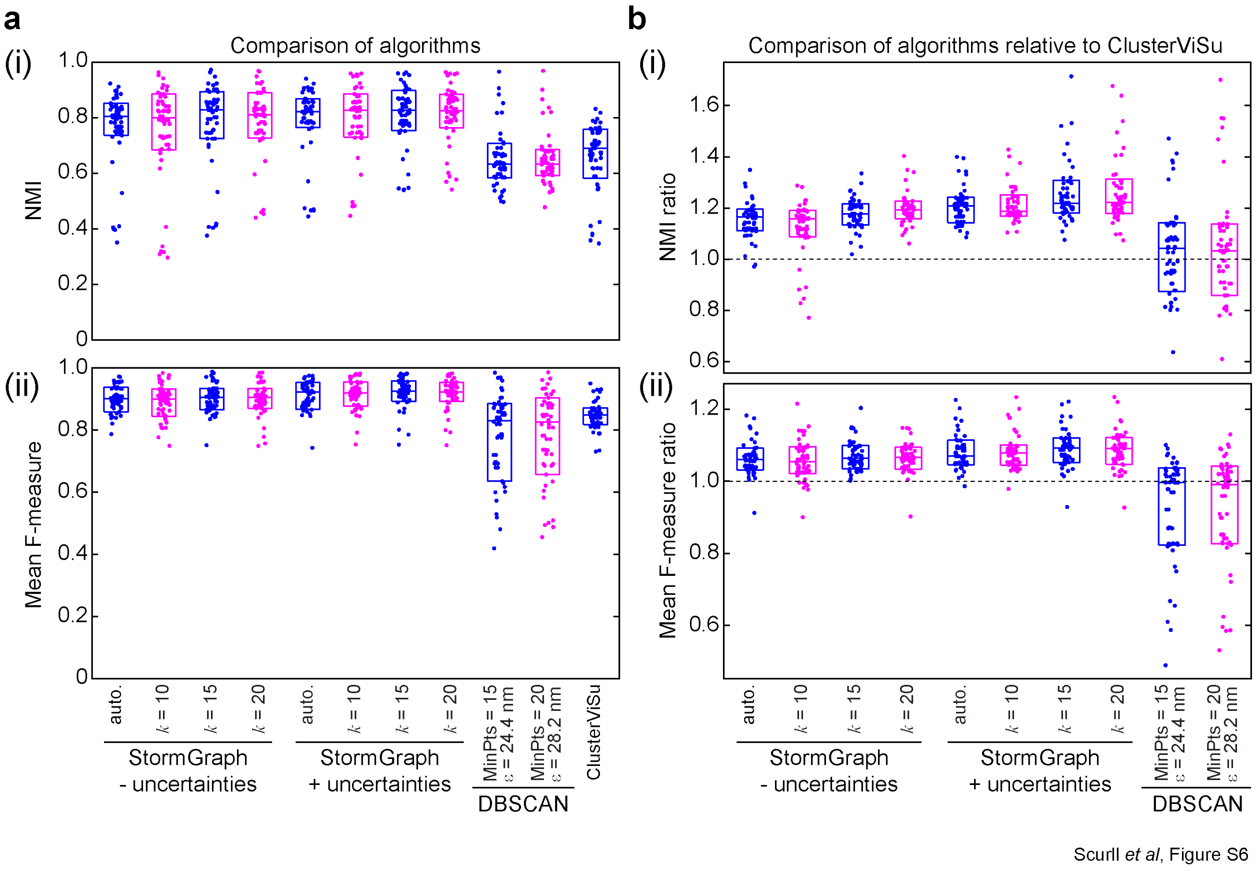


**Figure S6. StormGraph is more accurate than ClusterViSu and DBSCAN at assigning localizations to clusters in simulated data with no overcounting of single molecules.** We simulated 2D SMLM data as described for Figure 2 in the Methods but with every molecule having exactly one localization (i.e. no overcounting). We then performed the same algorithm testing that we performed in Figure 2. **(a)** Accuracy of cluster assignments by StormGraph, DBSCAN, and ClusterViSu, measured by (i) Normalized Mutual Information (NMI) and (ii) mean F-measure, for 38 simulated datasets in which every molecule is localized exactly once. StormGraph was implemented either using (+) or not using (-) localization uncertainties during clustering and with $r_0$ set either using the heuristic method (auto.) or using $k = 10$, 15 or 20. For all runs of StormGraph, $\alpha = 0.05$ and MinCluSize = 5 were fixed. DBSCAN was tested using 24 different pairs of its two user-specified parameters MinPts and $\varepsilon$. Shown here are the two choices of DBSCAN parameters that yielded the best results. A minimum cluster size of 5 points was used for ClusterViSu. ClusterViSu results are not shown for one of the 38 datasets because it failed to complete analysis in under 2 h. Boxes show medians and interquartile ranges. **(b)** Same as (a), but with StormGraph's or DBSCAN's accuracy, measured by (i) NMI and (ii) mean F-measure, divided by ClusterViSu's NMI or mean F-measure for each of the 37 simulated datasets for which ClusterViSu analysis completed in under 2 h. Ratios $> 1$ indicate that StormGraph or DBSCAN was more accurate than ClusterViSu for the corresponding datasets.

#### Figure S7

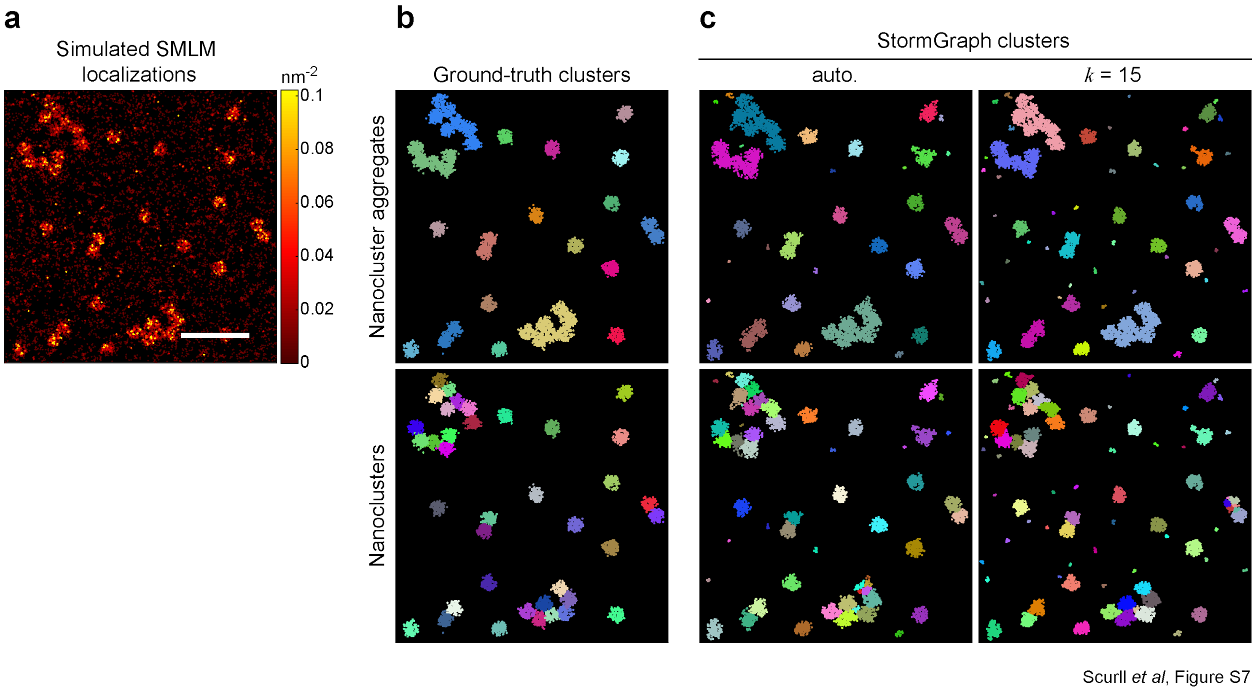


**Figure S7. StormGraph's cluster hierarchy accurately recovers clusters at different scales in simulated data. (a)** An example simulated SMLM dataset containing isolated and aggregated nanoclusters of radius 50 nm and randomly distributed molecules outside of clusters. This was one of the 64 simulated datasets used for Figure 2. Color bar = density (nm$^{-2}$), scale bar = 500 nm. For realism, simulated data included multiple localization of single molecules, which can lead to the appearance of small clusters of localizations among the non-clustered molecules. **(b)** Ground-truth clusters at the two levels of clustering in the simulated data from (a). Colors distinguish different clusters. Top: Large-scale clustering. Some nanoclusters are aggregated into larger clusters. Bottom: Individual nanoclusters (50 nm radius) at the small scale. **(c)** Clusters from the cluster hierarchy found by StormGraph ($\alpha = 0.05$, minimum cluster size = 5 localizations) using either the heuristic (auto.) method or the $k$NN method with $k = 15$ to set $r_0$ and using localization uncertainties. Top: Colors show distinct clusters at the top (coarsest) level of the cluster hierarchy. Bottom: Colors show distinct clusters at a manually identified level of the cluster hierarchy.

#### Figure S8

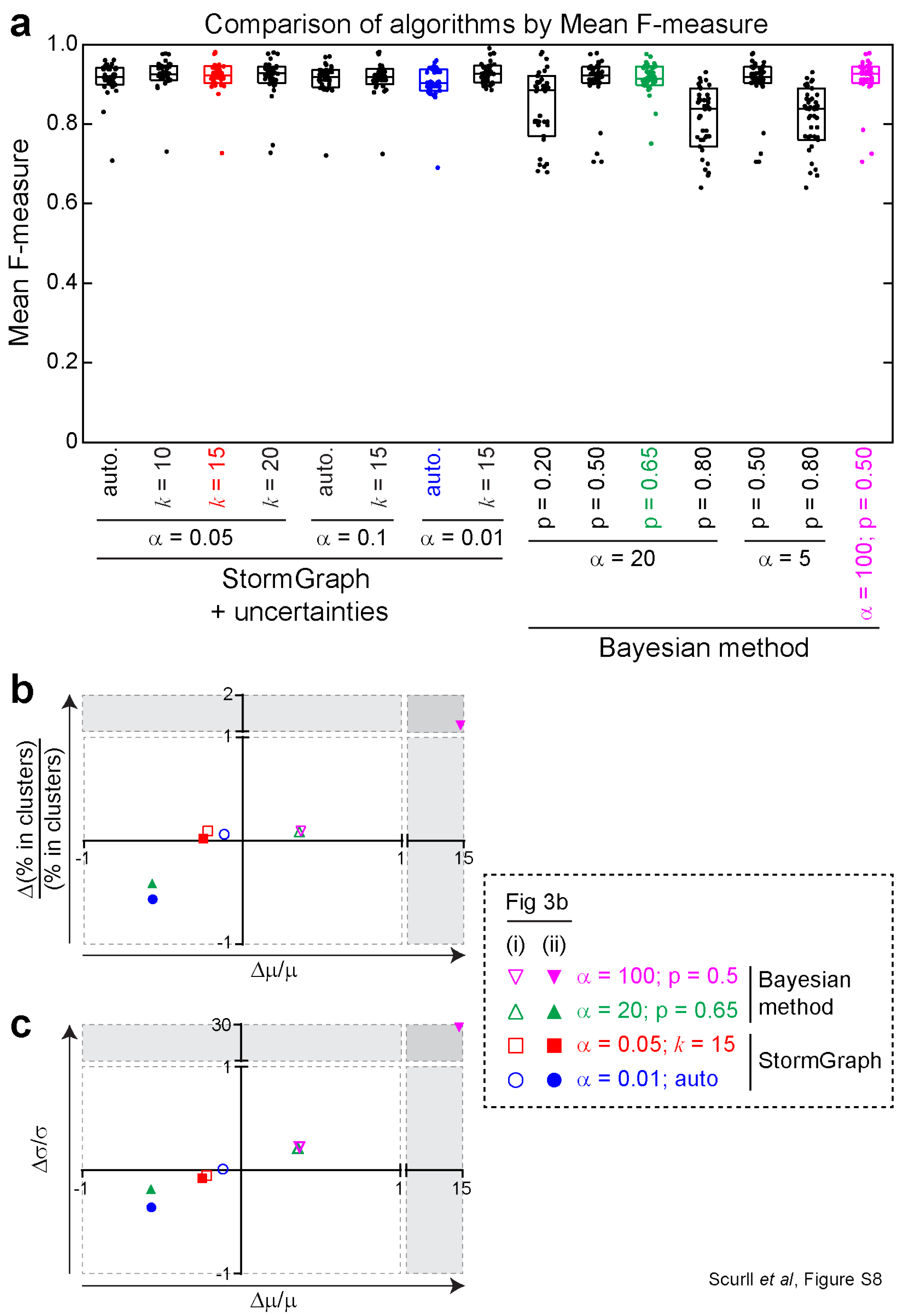


**Figure S8. Further comparison of StormGraph to the Bayesian clustering method. (a)** Mean F-measure values measuring the performances of StormGraph and the Bayesian method (10) at assigning localizations to clusters compared to ground truth (mean F-measure = 1 $\implies$ 100% match). For a variety of input parameter values, StormGraph and the Bayesian method were applied to 30 simulated 1 μm × 1 μm

ROIs. Each dot in the figure corresponds to one ROI. For StormGraph, localization uncertainties were always used (+ uncertainties), $r_0$ was set using either the kNN method or the heuristic (auto.) method, MinCluSize was not set, and the parameters $\alpha$ and $k$ were varied. The Bayesian method was always implemented using suitable fixed ranges of $R$ and $T$ values and the default prior distribution of cluster radii, which we expected to be suitable for our simulated data based on our simulation parameters, and the parameters $\alpha$ and $p$ were varied. **(b–c)** Cluster quantification errors by StormGraph and the Bayesian method relative to ground truth for the two example simulated ROIs shown in Figure 3b. The fractional error in the percentage of localizations assigned to clusters (b) and the fractional error in the standard deviation, $\sigma$, of number of localizations per cluster (c) are plotted *versus* the fractional error in the mean number of localizations per cluster, $\mu$, for each of the 30 simulated ROIs. N.b. $\pm 1$ means $\pm 100\%$ error, and errors greater than 100% are shown in the shaded regions, which have different axis scales. The parameter selections correspond to the highlighted parameters in (a–b).

#### Figure S9

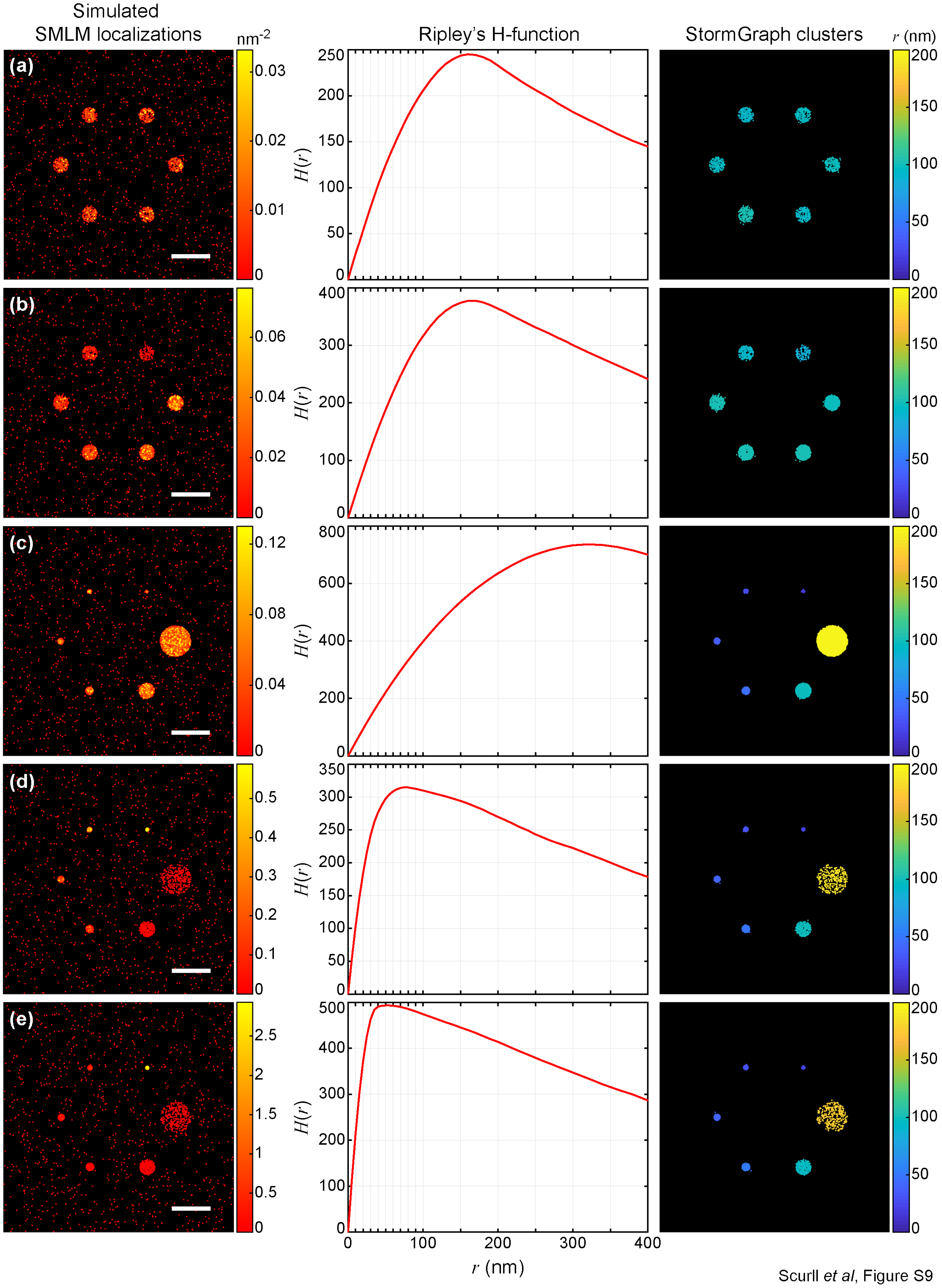

Simulated
SMLM localizations
nm-2
Ripley's H-function
StormGraph clusters
r (nm)
(a)
(b)
(c)
(d)
(e)
H(r)
r (nm)
0
100
200
300
400
0.03
0.02
0.01
0.06
0.04
0.02
0.12
0.08
0.04
0.5
0.4
0.3
0.2
0.1
2.5
2
1.5
1
0.5
250
200
150
100
50
400
300
800
600
350
500
Scurll et al, Figure S9

**Figure S9. The H-function derived from Ripley's K-function is unsuitable for quantifying SMLM data containing heterogeneous clusters.** Left panels: Six simulated circular clusters of various radii and densities, and approximately 1,000 randomly distributed points. Simulated data contained no uncertainties. Color bar = density ($nm^{-2}$), scale bar = 500 nm. Middle panels: Ripley's H-function vs length scale ($r$) for the simulated data. Deviation above 0 indicates clustering. The position ($r$ value) of the peak of $H(r)$ is often used to estimate a single value for cluster radius. Right panels: Clusters found by StormGraph and colored by their radii estimated from the StormGraph-quantified cluster areas (radius = $\sqrt{\text{area}/\pi}$). StormGraph was implemented using $k = 10$, $\alpha = 0.05$, and a minimum cluster size of 5 points. **(a)** All clusters contain approximately 200 points and have radius 100 nm. **(b)** Clusters all have radius 100 nm but contain different numbers of points (100, 200, 300, 400, 500, and 600). **(c)** Clusters all have the same density but have different radii (20 nm, 30 nm, 40 nm, 50 nm, 100 nm, and 200 nm). **(d)** Clusters all contain approximately 300 points but have different radii (20 nm, 30 nm, 40 nm, 50 nm, 100 nm, and 200 nm). **(e)** Smallest cluster (20 nm radius) contains approximately 1,000 points. All other clusters (radii 30 nm, 40 nm, 50 nm, 100 nm, and 200 nm) contain approximately 300 points.

### Figure S10

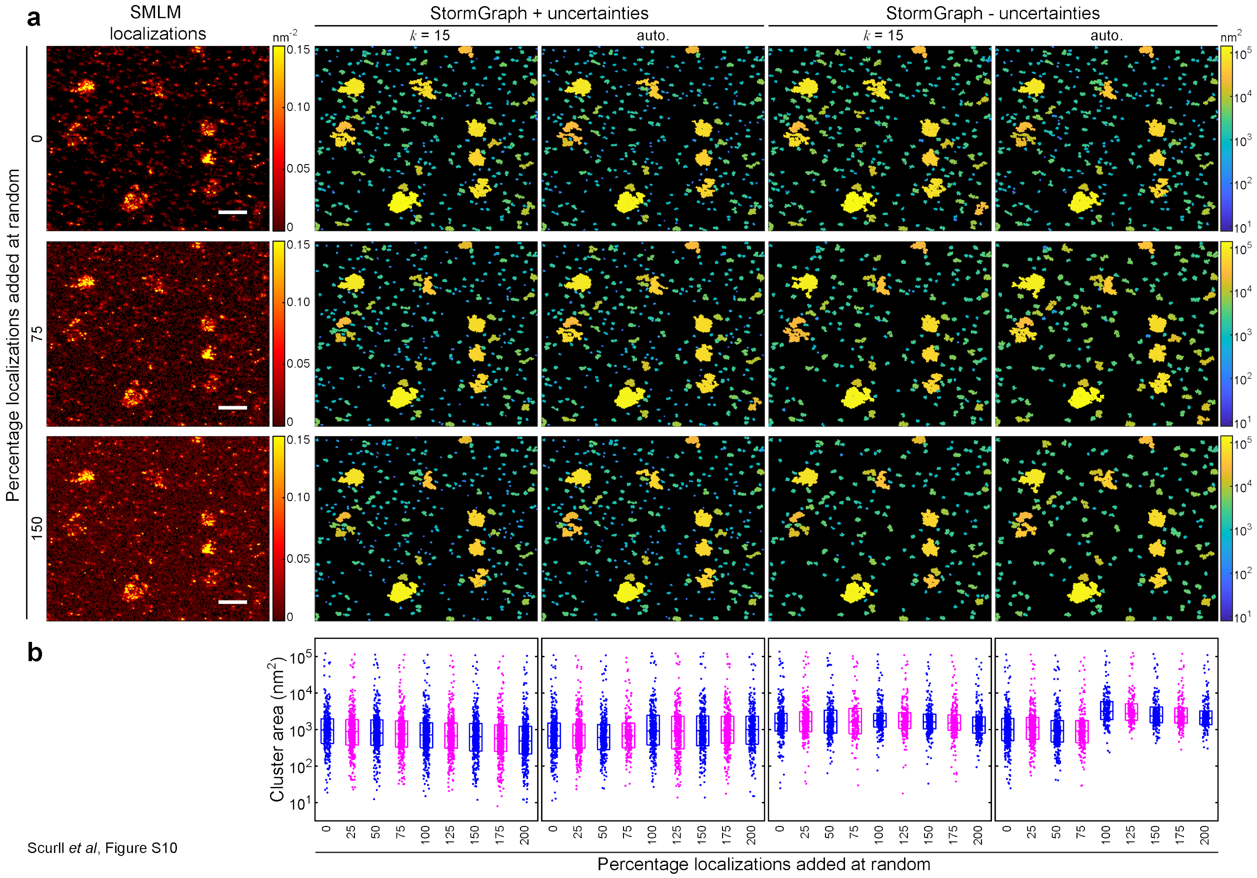


**Figure S10. Effect of increasing random noise on StormGraph analysis results. (a)** IgM-BCR dSTORM localizations in an ROI from an HBL-1 cell (top-left panel) with increasing numbers of additional randomly distributed points (left panels; color bars = density ($nm^{-2}$)). The number of random points added is specified as a percentage of the original number of localizations: 0% (top), 75% (middle) and 150% (bottom). For each case, distinct clusters were identified by StormGraph either using (+) or not using (-) localization uncertainties and using either the heuristic method (auto.) or k-nearest neighbor ($k = 15$) method to set $r_0$. Localization clusters possibly arising from overcounted single molecules were not removed. Color bars = cluster area ($nm^2$)). Scale bar = 500 nm. **(b)** Areas of all distinct clusters identified by StormGraph in the ROI when different numbers of randomly distributed points were added (up to 200% of the original number of localizations). Results are for the four implementations of StormGraph in the corresponding columns of panel (a).

#### Figure S11

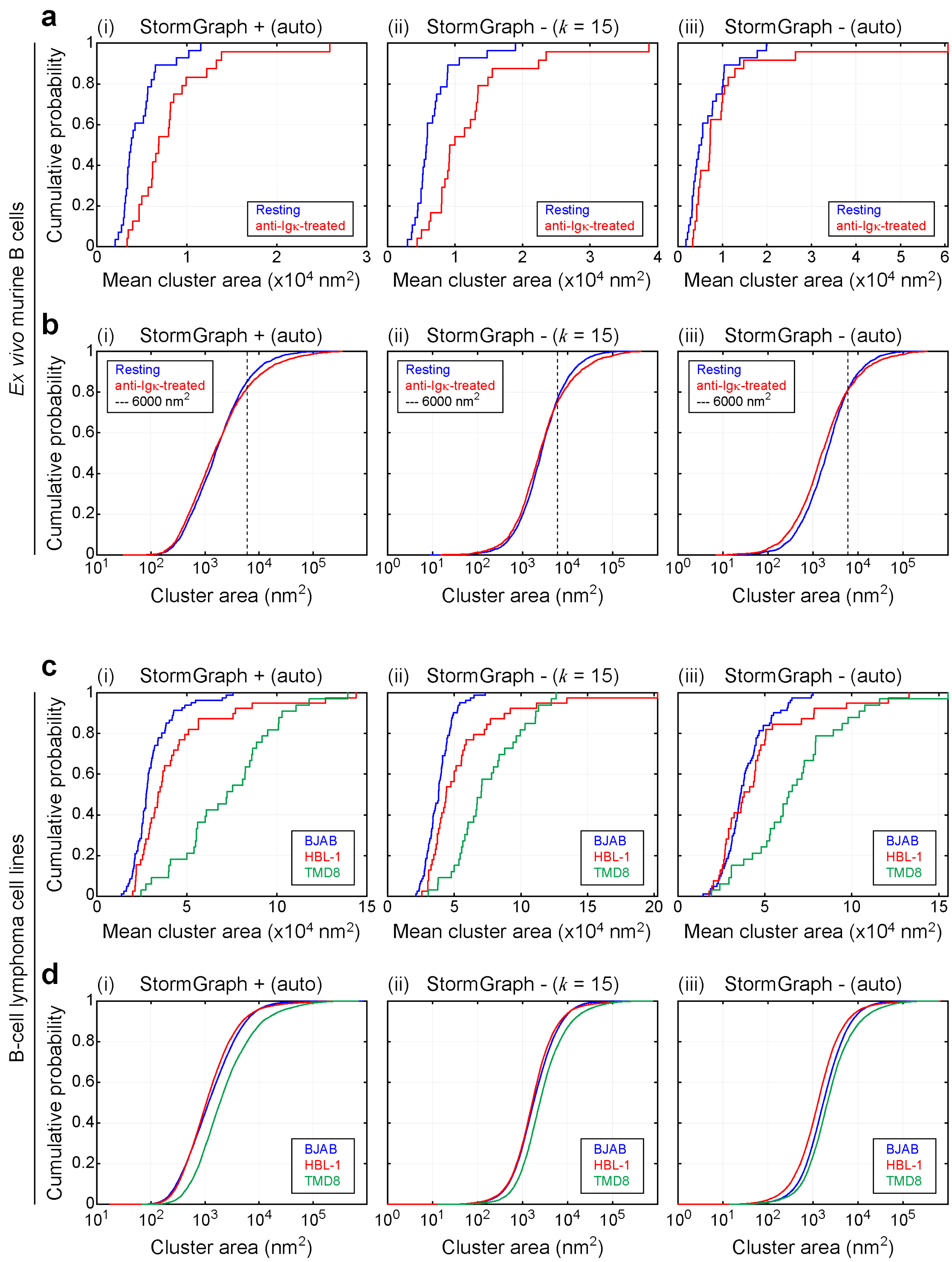


Scurll *et al*, Figure S11

**Figure S11. Assessment of how different StormGraph settings impact cluster analysis results for dSTORM data.** The dSTORM data for IgM-BCRs shown in Figure 5 was analyzed by StormGraph using either the heuristic method (auto.) or the kNN method with $k = 15$ to determine the value of $r_0$. Positional uncertainties in dSTORM localizations were either used (+) or not used (-) during analysis by StormGraph. **(a–b)** Cumulative distribution function (CDF) for (a) the mean area and (b) all areas of IgM-BCR clusters detected by StormGraph, using different settings, in dSTORM ROIs from resting (blue) and anti-Igκ-treated (red) *ex vivo* murine splenic B cells. **(c–d)** CDF for (c) the mean area and (d) all areas of IgM-BCR clusters detected by StormGraph, using different settings, in dSTORM ROIs from BJAB (blue), HBL-1 (red) and TMD8 (green) cells.

### Figure S12

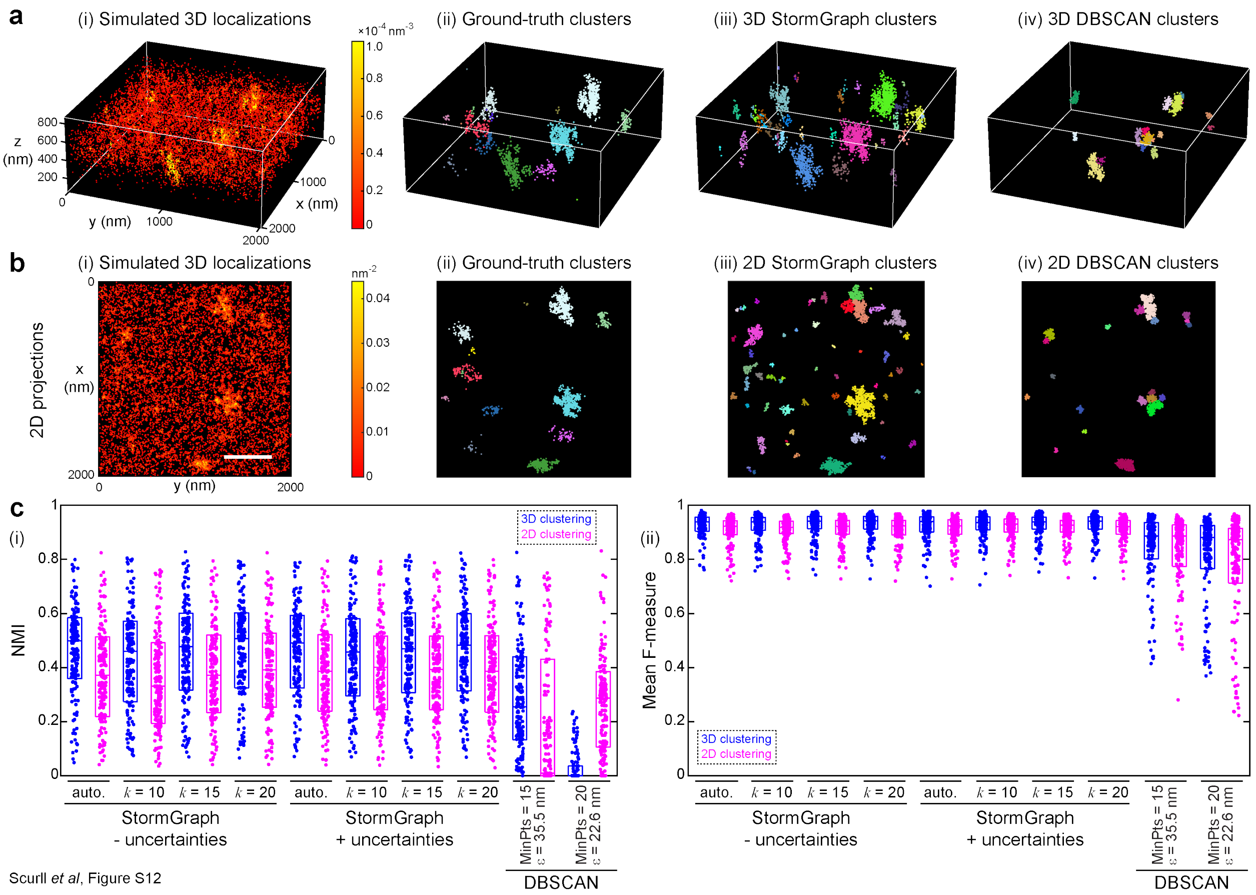


**Figure S12. StormGraph identifies clusters in 3D simulated data more accurately than DBSCAN. (a)** (i) Simulated 3D SMLM data example. Color bar = 3D density. In this example, the density of molecules outside of clusters was 5% of the within-cluster density of molecules, which still resulted in ∼85% of localizations not belonging to any true cluster. We also tested examples with up to four times this relative density of unclustered molecules. (ii) Ground-truth clusters in the simulated example. Colors distinguish distinct clusters. (iii) Clusters identified by StormGraph implemented in 3D with localization uncertainties used during clustering, $r_0$ determined heuristically (auto.), $\alpha = 0.05$, and MinCluSize = 5 localizations. Colors distinguish distinct clusters. (iv) Clusters identified by DBSCAN implemented in 3D using MinPts = 15 and $\varepsilon = 35.5$ nm. These were the DBSCAN parameter values that had the best average NMI scores for 3D clustering of 130 simulated datasets. **(b)** (i–ii) 2D projections of (a)(i)–(ii) respectively onto the $xy$-plane. (iii–iv) Clusters identified, in the 2D projection of the 3D simulated example, by 2D implementations of (iii) StormGraph, and (iv) DBSCAN. StormGraph was implemented in 2D using the same settings as the 3D implementation in (a)(iii). DBSCAN was implemented in 2D using MinPts = 20 and $\varepsilon = 22.6$ nm. These were the DBSCAN parameter values that had the best average NMI scores for 2D clustering of 130 simulated datasets. Color bar = 2D density; scale bar = 500 nm. **(c)** Accuracy of cluster assignments by 3D and 2D implementations (applied, respectively, to the full 3D data or to the 2D $xy$ projections) of StormGraph and DBSCAN for 130 simulated 3D datasets, as assessed by (i) normalized mutual information (NMI; 1 = perfect) and (ii) mean F-measure (1 = perfect). Boxes show medians and interquartile ranges. StormGraph was implemented using $\alpha = 0.05$ and MinCluSize = 5 localizations. Localization uncertainties were either used (+) or not used (-) during clustering, and $r_0$ was determined using the heuristic method (auto.) or using the indicated values of $k$. DBSCAN was implemented using the indicated parameters. The first set of DBSCAN parameters (MinPts = 15, $\varepsilon = 35.5$ nm) was optimized for 3D clustering and the second set (MinPts = 20, $\varepsilon = 22.6$ nm) was optimized for 2D clustering.

#### Figure S13

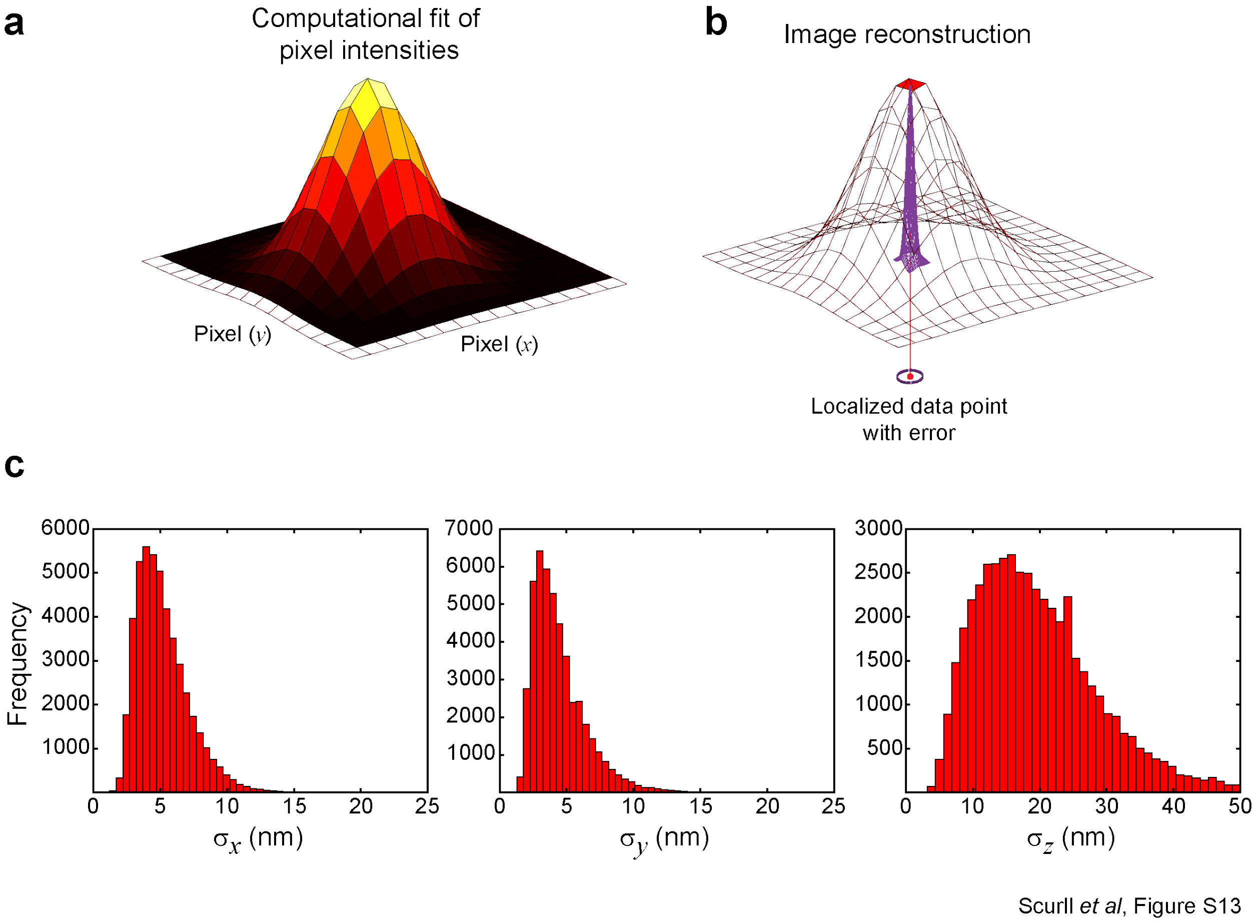


**Figure S13. Computational localization and positional uncertainty estimation of dSTORM fluorescence emission events. (a)** A point spread function (PSF) is fit to the fluorescence intensity profile of each SMLM blink. **(b)** The localization coordinates of each fluorescence emission event and their associated uncertainties (i.e. estimated measurement errors) are determined simultaneously from the computational fit of the PSF. **(c)** Distributions of the $x$, $y$ and $z$ positional uncertainties, expressed as standard deviations, in dSTORM localizations in a representative dataset.
